# Persistent viral infection in the *Drosophila* fat body is associated with immune activation at the single cell level

**DOI:** 10.64898/2026.01.06.697976

**Authors:** Nilanjan Roy, Robert L. Unckless

## Abstract

**Background:** Viruses are ubiquitous and can spread in two main ways: vertically, which involves transmission through or associated with gametes, and horizontally, which occurs through direct contact, airborne transmission, or indirect contact, such as through ingestion. Vertically transmitted, low virulence viruses can go undetected by both the immune system and researchers, and cause chronic, asymptomatic infections. In many *Drosophila* studies, researchers are unaware or ambivalent about the fact that the flies used may be infected with persistent viruses. Although they often have minimal or no observable fitness costs in laboratory fly samples, recent studies suggest that an increase in viral titer is associated with a decrease in lifespan.

**Results:** In this study, we explored changes in expression occurring during these cryptic virus infections. To achieve this, we utilized publicly accessible single-nuclear RNA sequencing (snRNA-seq) data of the *Drosophila* fat body, where we detected persistent infections of Nora and Drosophila A virus. We observed that Nora virus and Drosophila A virus exhibit broad cell-type tropism in the *Drosophila* fat body, and when coinfected, Drosophila A virus showed higher viral titer and cell infection rate. Transcriptomic analysis revealed substantial immune pathway alterations: Nora virus is broadly associated with upregulation of immune pathways (IMD, Toll), whereas Drosophila A virus is associated with downregulation of specific Toll pathway effector genes. Additionally, the expression of somatic transposable element (TE) transcripts was associated with viral infection, showing mating status-dependent patterns with downregulation in Nora virus-infected virgin flies and upregulation in mated flies for both viruses.

**Conclusions:** Overall, our results indicate that cryptic and persistent viral infections in *Drosophila* elicit transcriptional changes in the fat body, including activation of immune responses, and are associated with dysregulation of TE activity in somatic fat body cells.

## 1 Introduction

Organisms are often persistently infected with viruses, and these viruses have consequences for hosts [1, 2]. Nearly all humans acquire one or more herpesviruses in their lifetime [3]. Epstein–Barr virus (EBV, or human herpesvirus 4) infects up to 95% of adults globally [4]. Although EBV infection is typically asymptomatic, disruption of the virus-host balance can lead to serious outcomes, including cancer and autoimmune diseases [5]. Thus, understanding persistent virus-host interactions is crucial for identifying emerging diseases, monitoring their spread, and guiding public health responses [6, 7].

The fruit fly, *Drosophila melanogaster* has emerged as a valuable model organism for studying viral infections [8, 9]. A diverse range of RNA viruses infect *Drosophila*, including Nora virus and Drosophila A virus, which belong to the order *Picornavirales* and are evolutionarily related to vertebrate picorna viruses [10, 11, 12, 13, 14, 15]. The innate immune response pathways in *Drosophila* also share many similarities with those of vertebrates [16, 17]. Toll signaling in *Drosophila* corresponds to Toll-like receptor (TLR) signaling, IMD signaling represents TNF*α* receptor and cGAS/STING signaling [18]. RNAi signaling in *Drosophila* reflects both RNAi and RIG-like receptor signaling in vertebrates. RIG-like receptors (which detect viral dsRNA) and Dicer (a major component of RNAi, cleaves viral dsRNA) share similar superfamily 2 (SF2) helicase domains which suggests that this helicase domain originally involved in an antiviral role in a common ancestor [19]. Similarly, JAK-STAT signaling is conserved between *Drosophila* and many vertebrates [18].

The most studied persistent viral pathogen in *Drosophila* flies is the *Drosophila* C virus (DCV) [20]. It is a positive sense RNA virus which enters into the cells by clathrin-mediated endocytosis. Depending on the viral dose and genetic makeup, DCV infection results in full mortality in 3–13 days after infection [21, 22, 23, 24]. However, a significant gap in knowledge exists regarding other persistent viral infections in *Drosophila*, particularly for Drosophila A virus and Nora virus. These viruses naturally infect *Drosophila* in the wild and are found in *Drosophila* laboratory lines [25].

Nora virus (single-stranded positive-sense RNA virus) has a genome size of approximately ∼12.3kb and encodes four open reading frames (ORFs) [12]. ORF1 of Nora virus functions as an RNAi suppressor in *Drosophila melanogaster* [26]. The virus is typically shed in feces, enabling transmission from mother to offspring through contaminated food or the environment [12], which facilitates its persistent presence in laboratory lines. Infections can vary greatly in viral load among individuals within the same population and usually do not cause overt symptoms. However, Nora virus has been associated with reduced lifespan, abnormal locomotor activity, and increased susceptibility to secondary bacterial infections [13, 27, 28, 28]. Studies in the gut have shown induction of genes involved in the Toll, immune deficiency (IMD), and cGAS-sting pathways, suggesting gut-specific immune response [29, 30].

Drosophila A virus (single-stranded positive-sense RNA virus) genome size is approximately ∼4.5kb and has two ORFs [31]. While it remains less studied than Nora virus, recent work has shown that Drosophila A virus triggers the cGAS/Sting pathway in the *Drosophila* gut [32]. Like Nora virus, it causes a cryptic infection that can lead to reduced lifespan, decreased reproductive output, and the activation of immune responses [32].

Previous studies of Drosophila A virus and Nora virus have primarily focused on cell culture models or whole-tissue analysis or in the gut (primary infection site), leaving gaps in our understanding of host-pathogen interactions at the cellular level in tissues. In particular, persistent viral infections have rarely been studied in the *Drosophila* fat body—the primary immune organ—despite its central role in antiviral defense [33].

Host responses to viral infections are most often examined through the lens of protein-coding gene expression. However, our understanding of how viral infections influence the dark matter of the genome, particularly transposable elements (TEs) remains limited. Viruses and TEs have a complex relationship both in terms of their genetic structure and also the host reponse against them [34, 35, 36]. TEs can move around in the genome, causing genome rearrangements (in germline cells) and differences in transcription [37, 38]. Studies have shown that viral infections can disrupt the expression of TEs, raising important questions about whether such changes benefit the virus, the TEs, or both [39, 40, 41]. TEs, such as retrotransposons, display sequence similarities to viruses. Retrotransposons originated from ancient viral infections that became integrated into host genomes [42]. A classic example is the *Ty3* retroelement (formerly *gypsy*) in *Drosophila melanogaster*, which is widely distributed among drosophilids and encodes a retroviral-like *env* gene that produces a functional envelope protein, thereby conferring infectious capability [43, 44, 45]. In germline cells and somatic cells within the gonad, the piRNA pathway (small RNA system) suppresses TEs to protect genome integrity [46]. The RNA interference (RNAi) pathway, another small RNA system, serves as a key antiviral defense mechanism [47]. Moreover, RNAi is a very well validated regulator of somatic TE expression[48]. This suggests a potential evolutionary link between the small RNAs that protect against viral infections and those that defend against TEs, reflecting shared origins and blurred boundaries between immunity against viruses and TEs.

Despite significant advances in understanding host-virus interactions using *Drosophila*, most studies have relied on bulk RNA-sequencing (RNA-seq), which measures average gene expression across all cells in a tissue [49]. While this approach provides valuable insights, it masks expression heterogeneity among distinct cell types [50]. This limitation is particularly important in the context of viral infections, where diverse cellular responses can shape the course and outcome of infection [51, 52]. Single-nuclear RNA-sequencing (snRNA-seq) offers a powerful alternative by enabling high-resolution analysis of gene expression at the individual cell level, thus uncovering the complexity of transcriptomic responses within cell types in tissues [53, 54, 55, 56]. In this study, we used publicly available snRNA-seq data to investigate the cell type tropism, host responses, and regulation of TEs in Nora and Drosophila A virus infection in the *Drosphila* fat body.

## 2 Materials and Methods

### 2.1 Dataset structure

To study transcriptional changes associated with Nora and Drosophila A virus infection at single nuclear level, we utilized *Drosophila* data from a study of fat body tissue [57] as the fat body is the main immune organ of *Drosophila* [58, 59, 60]. We leveraged publicly available *Drosophila* fat body single-nuclear RNA (snRNA) sequencing data from NCBI (Bioproject: PRJNA698971). The experiments were performed on wildtype *Drosophila melanogaster* Canton S flies. Details of the fly rearing and mating protocols along with all other procedures are described in the original article [57]. The experiments used bacterial infection to demonstrate the fat body’s role in balancing its immune and metabolic functions after mating. Virgin and mated females flies were infected with the gram-negative bacteria *Providencia rettgeri*. We only used virgin uninfected and mated uninfected replicates for our subsequent analysis to avoid complications from experimental bacterial infections in the original study. The snRNA sequencing experiment had two replicates over four treatments: virgin uninfected, virgin infected, mated uninfected, and mated infected. Two full blocks of the experiment were conducted on separate days two months apart. Briefly, nuclei were suspended from the fat body in PBS containing 2% BSA for 10X chromium-based RNA sequencing. RNA libraries were made with 10X chromium v3 chemistry. Libraries were sequenced on the Illumina platform at 10X recommended conditions.

Through a virus screening pipeline (described below), we detected Nora and Drosophila A virus reads in specific snRNA sequencing samples in different treatment groups (Supplementary figure S1). To confirm that putative viral reads were not mismapped from host genes, we performed BLASTN searches of viral sequences against *Drosophila* genes [61].

We found no BLASTN hit of either Nora or Drosophila A virus sequence against *Drosophila* genes even with a relatively low E value cutoff of 1 × 10*^−^*^5^. Viral reads were also mapped to the host genome and they showed no alignment. These findings allowed us to study Nora and Drosophila A virus infection dynamics and transcriptional changes in the host due to infection.

### 2.2 Identification of viruses in the fat body snRNA-Seq data

To determine what viruses might be infecting the samples, we pulled all the known *Drosophila* virus sequences (fasta file) from Obbard lab website (obbard.bio.ed.ac.uk/data/Updated_Drosophila_Viruses.fas.gz) [25, 62, 63]. We then used BWA mem (version 0.7.18-r1243-dirty) to align the fat body snRNA sequencing reads to these viral sequences [64]. Samtools coverage (version 1.17) was used to quantify viral reads [65]. We employed a hard threshold of 100 RPM (reads per million) to determine whether the sample was considered infected with virus. That means if a viral sequence had at least 100 RPM, we considered that sample to be infected with that specific virus. While this does not guarantee a robust infection, it reduces the likelihood of contamination or passing association. This approach enabled the identification of viruses present in the snRNA-seq *Drosophila* fat body samples.

Later, we mapped snRNA-seq reads to a combined reference genome that included *D. melanogaster* transcripts (BDGP Release 6 + ISO1 MT/dm6) and viral sequences that passed the aforementioned threshold. This allowed us to analyze transcriptional changes associated with viral infection in the *Drosophila* fat body (see details below).

### 2.3 Data processing and analyses

Barcode processing along with count matrix generation was performed separately for each replicate of the virgin uninfected and mated uninfected treatment separately using CellRanger (version 7.2.0) [66]. We used the filtered gene-barcode matrix generated by CellRanger to omit barcodes associated with background noise (likely from free floating, ambient mRNA from lysed or dead cells). For mapping snRNA sequencing reads, We used the *Drosophila melanogaster* reference genome (BDGP Release 6 + ISO1 MT/dm6) from UCSC genome browser (genome.ucsc.edu). We modified the transcriptome files (fasta and gtf) by adding Nora virus (accession JX220408.1), and Drosophila A virus (accession KP969946.1) sequences. The count matrix was processed and analyzed with Seurat (version 5.0.0). For creating the seurat object, we used minimum cell count of 3 and minimum features of 200 threshold. We merged each replicate of the virgin uninfected and mated uninfected treatment to do combined analyses. We used Harmony (version 0.1.1) for batch correction and integration of replicates of the virgin uninfected and mated uninfected treatments [67]. Cells containing between 200 and 2500 genes and *<*25% mitochondrial genes were retained. By doing so, we remove low-quality cells or empty droplets, which often have very few genes, as well as cell doublets with an abnormally high gene count. Scaling and log normalization were done with the standard seurat functions with default parameters. The top 50 PCs (based on variance; Elbow plot) were used for Principal component analysis (PCA) and Uniform Manifold Approximation and Projection (UMAP). Cell types (clusters) were generated at a 0.1 resolution. We intentionally kept the clustering resolution low as this helped us to separate canonical fat body cell types from the other non-fat body cell types. The *Drosophila* fat body is difficult to dissect and as a result, there is likely contamination of cell types from other nearby tissues [68]. Cell types were identified using marker genes with Cell Marker Enrichment tool from Flyrnai (flyrnai.org/tools/single_cell/web/enrichment). We used sc-Customize (version 2.1.2) to calculate the percentage of virus infected cells in different cell types [69].

Differential gene expression analyses were performed with the analysis of covariance (ANCOVA) method as this allowed to control for covariates such as cell types. The R package Companion to Applied Regression (car, version 3.1-2) was used for ANCOVA. Before doing the differentially expressed genes analyses, we defined cells as virus infected if the cell had viral titer (normalized count) greater than 0.

To identify differentially expressed genes and to determine which genes showed cell type–dependent effects of Nora virus (NV) infection in virgin flies, we used the following model, where 0 (categorical value) indicates virus-uninfected cells and 1 (categorical value) indicates virus-infected cells:

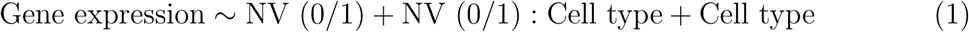

For mated flies, which were infected with both Nora virus and Drosophila A virus, the models were modified accordingly. We included the replicate covariate in the model here because Drosophila A virus was found in mated uninfected replicate 1, while both Drosophila A virus and Nora virus were present in replicate 2. So, to determine differentially expressed genes and to determine which genes showed cell type–dependent effects of Nora (NV) and Drosophila A virus (DAV) infection in mated flies we used the following model:

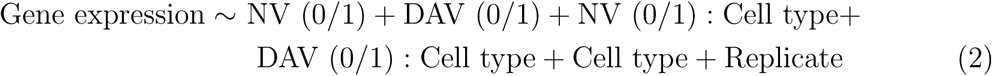

Pathway enrichment analysis was conducted with Flyenricher [70, 71]. SoloTE (version 1.09) was used to map the TE transcripts [72]. BAM files generated by CellRanger for selected samples (virgin uninfected and mated uninfected) were utilized as input for SoloTE analysis. CellRanger assigns reads to annotated genes only if reads map within one of their exons; thus, SoloTE initially removes gene-assigned reads to prevent misidentifying gene-contained TEs as expressed. SoloTE then evaluates remaining unassigned reads for overlaps with TE annotations and quantifies TE expression at both locus and family levels. We used repeatmasker output from BDGP Release 6 (dm6) for TE annotation. We derived the mean counts of TEs, piRNA pathway genes, and RNAi pathway genes from the SoloTE output. The mean count of TEs was calculated by including all the cells in the fat body tissue. We could not perform cell type specific analysis because the expression of piRNA pathway genes in the fat body is generally very low (13% of the total cells in the fat body had piRNA pathway genes expression). For this reason, mean counts of piRNA pathway genes were calculated from only cells that had piRNA pathway genes expression. However, we downsampled the total number of cells before calculating mean counts of piRNA pathway genes. It was done to randomly select cells that expressed piRNA pathway genes. R (version 4.3.1) and R packages tidyverse (version 2.0.0), pheatmap (version 1.0.12) were used for data wrangling and plotting. Additional figure editing was done using Microsoft PowerPoint.

## 3 Results

### 3.1 Nora and Drosophila A virus are not associated with specific fat body cell types

As the primary organ of innate immune defense in *Drosophila*, the fat body serves as a key tissue for investigating viral infections in *Drosophila* [33]. Yet, we have limited understanding of viral tropism in the fat body. We used cryptic Nora and Drosophila A virus infection to investigate tropism by examining viral RNA reads across different fat body cell types.

We found Nora and Drosophila A virus RNA reads in the fat body snRNA sequencing data (Gupta *et al.* 2022, [57]) but not consistently among treatments or replicates (Figure 1a, b). In replicate 1 of the virgin uninfected flies we did not find any viruses and in replicate 2 of the virgin uninfected flies, we only found Nora virus. In replicate 1 of the mated uninfected flies, only Nora virus RNA reads were detected, whereas replicate 2 contained reads from both Nora and Drosophila A virus (Figure 1a, b).

**Figure 1:**
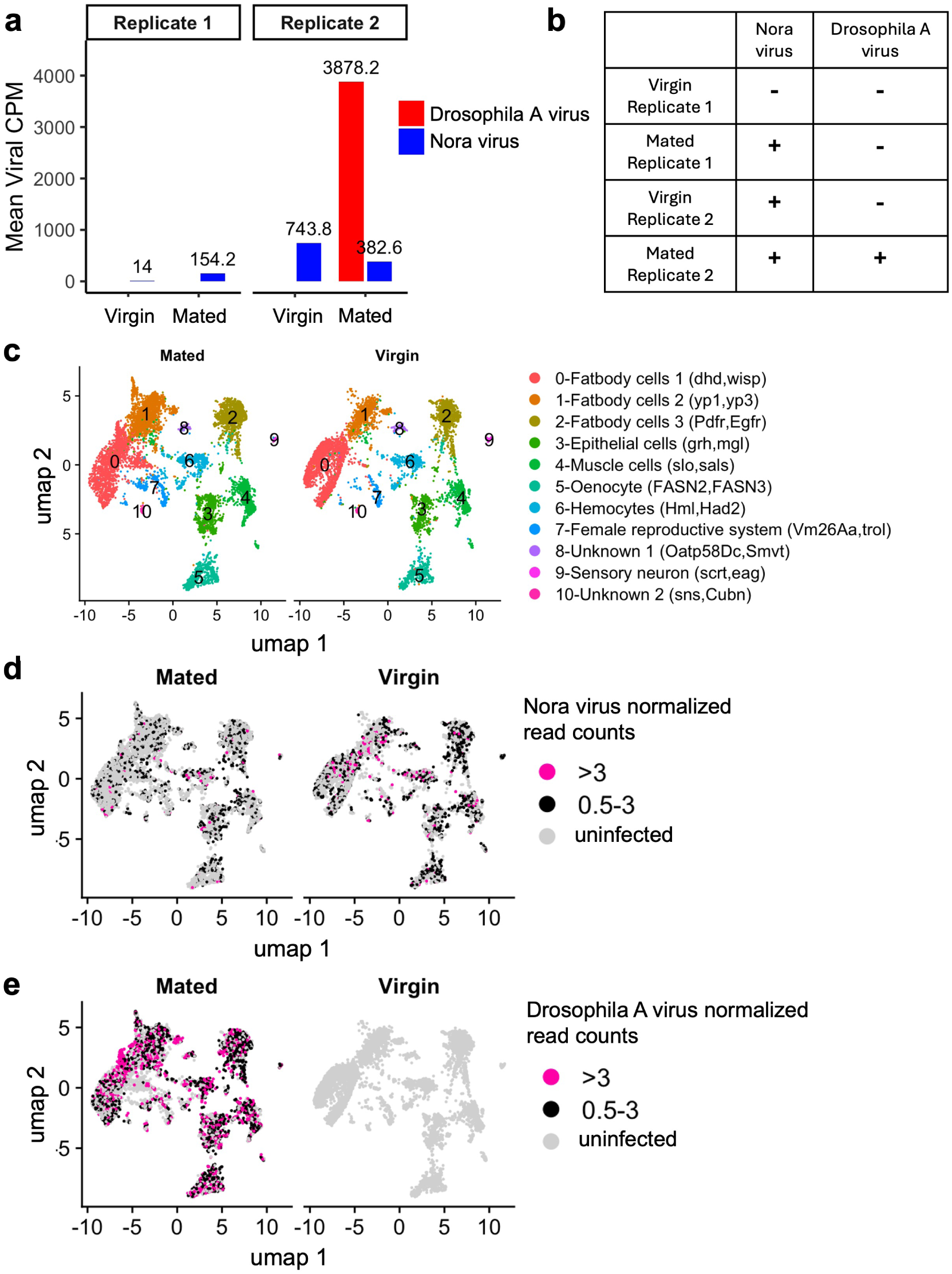
Nora and Drosophila A virus tropism in fat body cell types. (a) Nora and Drosophila A virus reads (CPM, counts per million) in different replicates of the snRNA-sequencing experiment in the fat body. (b) Viral infection status in each replicates in virgin and mated flies. (c) UMAP plot of 11 fat body-associated clusters. Colors and numbers in the UMAP indicate respective inferred cell types (d) Nora virus tropism in different fat body cell types in mated and virgin flies. (e) Drosophila A virus tropism in different fat body cell types in mated and virgin flies. Read counts are log normalized. Seurat LogNormalize method was used where feature counts for each cell are divided by the total counts of that cell, multiplied by a scale factor of 10,000, and then taking natural log–transformed value using log1p. *>*3 represents cells where viral read counts are greater than 3. 0.5-3 represents cells where viral read counts are between 0.5 to 3. Uninfected represents cells that don’t have any viral RNA. Log normalized counts can be converted to CPM using the formula *cpm* = (*e*^LogNormalized^ ^count^−1)×100. Thus, values *>* 3 correspond to viral RNA reads greater than ∼2000 CPM, while values between 0.5 and 3 correspond to viral RNA reads between ∼65–2000 CPM.

Viral reads mapped to various open reading frames (ORFs), with the majority aligning to capsid protein regions for both Nora and Drosophila A virus (Supplementary figure S2). Although these reads span the entire viral genome, the uneven coverage suggests the presence of both genomic and subgenomic RNAs. It is not uncommon that viral RNA reads are mapped to specific ORFs more than other ORFs in RNA sequencing based experiments [73, 74, 75].

After processing the snRNA sequencing fat body data, we found 11 clusters (Figure 1c). Clustering was performed with all the cells, then split by mating condition. Among them, fatbody cells 1 (*dhd, wisp*), fatbody cells 2 (*yp1, yp3*), and fatbody cells 3 (*Pdfr, Egfr*) are the three main fat body cell clusters. Details of the marker genes of each fat body clusters (along with GO enrichment terms) are in supplementary tables ST1, ST2. As described earlier, we used marker genes from Gupta *et al.* 2022 [57] to identify different clusters in the fat body. As the fat body is hard to dissect, there may be contamination from other nearby tissue. Perhaps as a result, we found 8 other non fat body cells types (epithelial cells, muscle cells, oenocyte, hemocytes, female reproductive system, unknown 1, sensory neuron, unknown 2). Gupta *et al.* 2022 [57] identified 18 distinct cell clusters in the *Drosophila* fat body. In our reanalysis of the dataset, we identified 11 clusters, 9 of which correspond to clusters reported in their study. Cell type clusters of all replicates of virgin and mated flies used in this study is in the Supplementary figure S3. In particular, both analyses consistently identified 3 main fat body cell clusters. The non-fat body cells can be contamination from nearby tissues or are rare, intrinsic components of the fat body.

To determine Nora and Drosophila A virus tropism, normalized viral RNA was used as a proxy for viral titer (Figure 1d, e). If a cell had normalized viral RNA count greater than 0, we classified the cell as infected. Based on this criteria, we found Nora virus RNA in all cell types in mated and virgin flies (Figure 1d). We explored alternative thresholds for determining infection status by using viral RNA cutoffs of *>*0.5, *>*1, and *>*1.5 instead of the initial *>*0 threshold. Consistent results were observed across all viral RNA cutoffs (Supplementary figure S4). This may indicate that Nora virus can indeed infect different cell types associated with the fat body. Drosophila A virus RNA was only found in the mated flies but also in all fat body and other cell types regardless of threshold for classifying cells as infected (Figure 1e, Supplementary figure S5). Thus, neither virus showed strong cell-type tropism in the fat body.

### 3.2 Drosophila A virus exhibits a higher viral titer and infects a greater number of cells compared to Nora virus

Although viral RNA was detected in multiple fat body cell types, its abundance may vary among them. We therefore sought to determine whether certain cell types are more heavily infected than others.

To identify which cell types are most likely to be infected, We fitted binomial logistic regression models separately for Drosophila A virus and Nora virus. The response variable was the number of infected versus uninfected cells, and the predictors included cell type, replicate, and the infection status of the other virus (e.g., Nora virus status when modeling Drosophila A virus infection, and vice versa). In the virgin flies, where Drosophila A virus was absent, we excluded its infection status from the model when assessing Nora virus infection. We then performed pairwise comparisons between cell types using Tukey’s Honest Significant Difference (HSD) test to identify specific cell type pairs with significantly different infection probabilities. We found that for both *Drosophila* A and Nora virus, fat body cell type 3 exhibited the highest likelihood of infection among the three major fat body cell types, with statistically significant differences observed in both virgin and mated flies (p-value *<* 0.01, Supplementary figure S6, Supplementary table ST3). Additionally, we observed a significant positive correlation between the proportion of cells infected by one virus and the proportion infected by the other across cell types (p-value *<* 0.001, Supplementary figure S7, Supplementary table ST3). This suggests that while there was no absolute cell type tropism for the viruses, some cell types (in particular fat body cells 3) are more prone to infection than others.

In addition to the proportion of cells infected, we noticed variation in the relative viral titer (as measured by the normalized read count). To investigate the variation in Nora and Drosophila A virus titers, we analyzed (ANOVA followed by post-hoc Tukey test) the number of normalized viral RNA counts in each fat body cell type. We performed the analysis in two ways. Either with only cells that harbored virus (viral RNA reads *>*0) (Figure 2b), or with all the cells. In both approaches, in the mated flies where coinfection was observed, the viral titer of Drosophila A virus was significantly higher (pvalue *<* 0.00001) across all cell types than Nora virus (Figure 2b, Supplementary figures S8, S9). We observed that Nora virus viral titer was higher in virgin flies than mated (pvalue *<* 0.0001, Supplementary figures S10, S11). A similar pattern was also seen in the comparison between virgin replicate 2 and mated replicate 1 (1a). Fat body cell type 3 exhibited significantly lower viral titers relative to the other two fat body cell types for both virgin and mated flies, for both viruses in virus expressing cells (p-value *<* 0.000001, Figure 2b, Supplementary table ST4). For example, Nora virus in the virgin flies, the mean viral titer of fat body cell 3 was lower than that of fat body cell 1 (mean difference of normalized viral RNA reads: fat body cell 1 minus fat body cell 3 = – 0.93, p-value *<* 0.000001), whereas the difference between fat body cells 2 and 1 was not significant (p-value = 0.865).

**Figure 2:**
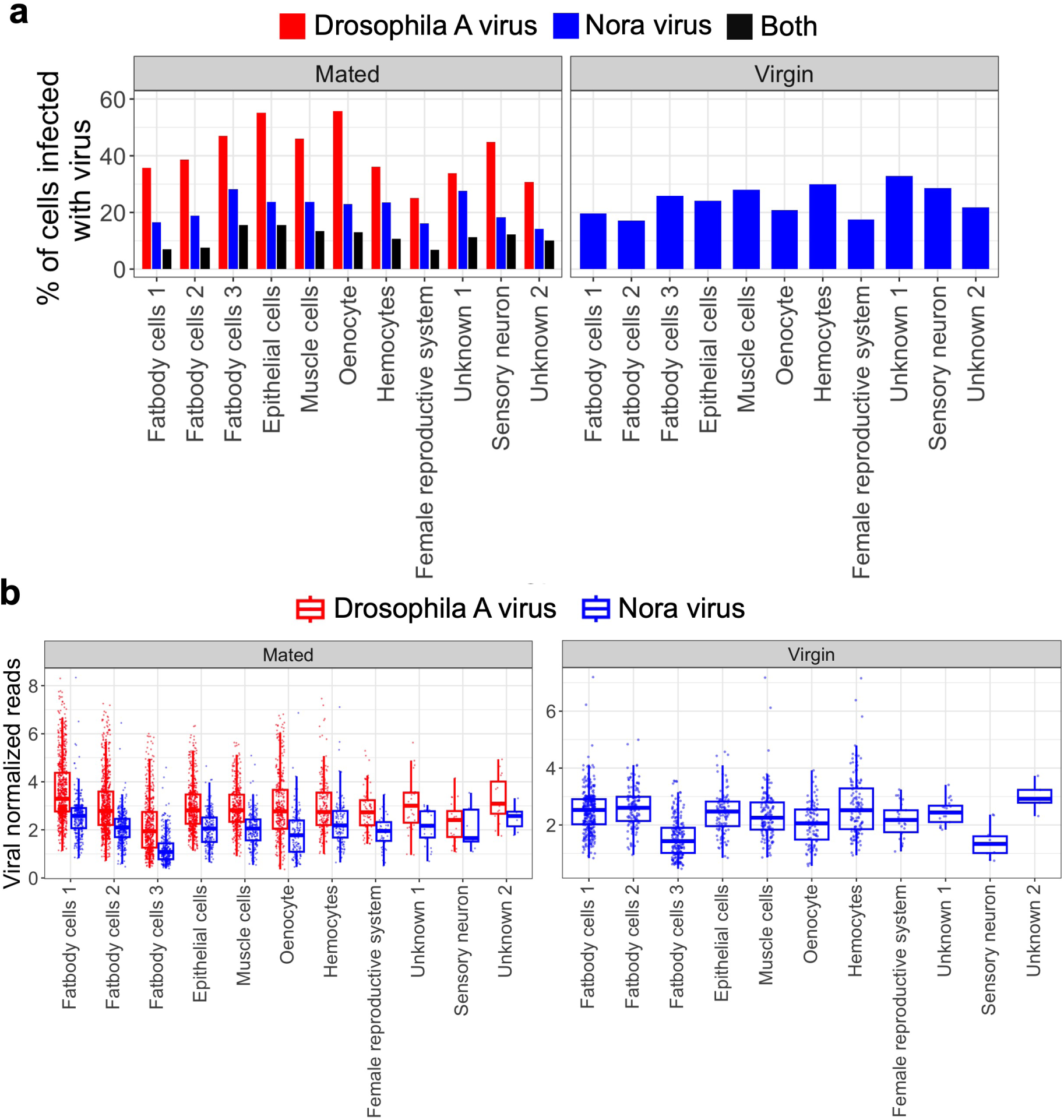
Infection profiles of Nora and Drosophila A virus in the fat body. (a) Percentage of infected cells in mated and virgin flies in Nora and Drosophila A virus infection. The percentage of virus-infected cells was determined by identifying cells with viral RNA read counts greater than zero. Both indicates cells containing RNA from both Nora and Drosophila A viruses. (b) Viral normalized read counts in mated and virgin flies in Nora and Drosophila A virus infection. Viral reads are normalized using Seurat LogNormalize method where feature counts for each cell are divided by the total counts of that cell, multiplied by a scale factor of 10,000, and then taking natural log–transformed value using log1p. Log normalized counts can be converted to CPM (counts per million) using the formula CPM= (*e*^LogNormalized^ ^count^ − 1) × 100. So, for example, log normalized value of 3 corresponds to viral RNA reads of ∼2000 CPM. Each dot represents an infected cell with viral RNA counts greater than zero.

A negative association (although statistically not significant) was observed between the infection rate (percentage of cells infected with the virus) and viral titer (viral RNA) for both Nora and Drosophila A virus in fat body cells (Supplementary figure S12). This observation may be attributed to the fact that susceptibility to infection (how easily a virus can enter a cell) and permissiveness to replication (how well a virus can replicate within a cell) are distinct properties, each influenced by different cellular factors. Alternatively, it could be that susceptible cells die. However, we hesitate to speculate further since this relationship was not significant.

In conclusion, Drosophila A virus may infect more cells in the fat body compared to Nora virus. Fat body cells 3 may be more prone to infection among the three main fat body cells and viral titer may be negatively associated with number of infected cells.

### 3.3 Expression of IMD and Toll pathway genes are altered in Nora and Drosophila A virus infections

The effects of viral infection on gene expression in the fat body remain poorly characterized. To address this, we performed a differential gene expression analysis in Nora and Drosophila A virus infection to understand infection-associated changes in the fat body. This is the main effect of virus infection. The design of the analysis is illustrated in Supplementary figure S29.

We first did the analysis on the virgin flies. There were two replicates of the virgin flies. Replicate 2 had Nora virus infection. As described earlier there is no Drosophila A virus infection in this condition. To get the differentially expressed genes due to Nora virus in virgins, we applied two strategies. In strategy 1, we compared the gene expression profile of replicate 2 (where Nora viral reads in cells is *>*1) with that of replicate 1 (where Nora viral reads in cells is 0). Figure 3a shows that, IMD *(DptA, AttC)* and Toll pathway *(Drs, Mtk)* genes are upregulated in association with Nora virus. IMD and Toll pathway are the two major innate immunity pathways in *Drosophila*. The IMD and Toll pathways have traditionally been associated with immune responses against fungal and bacterial infections, whereas the cGAS/STING and RNAi pathways are primarily linked to antiviral defense. However, recent studies have increasingly demonstrated that the IMD and Toll pathways also play significant roles in antiviral immunity [76, 77, 78]. Together with our analyses, these findings suggest that the IMD and Toll pathways may be critical for modulating immune responses to viral infections, or alternatively, that viral infection triggers a broad immune activation encompassing these canonical pathways. In addition to innate humoral immunity, muscle contraction genes such as Mlc2 (Myosin light chain 2), Mhc (Myosin heavy chain) are also up-regulated in association with Nora virus infection. Geotaxis defects and locomotor abnormality are some phenotypes associated with Nora virus infection in files [27], which might be associated with these upregulated muscle contraction related genes.

**Figure 3:**
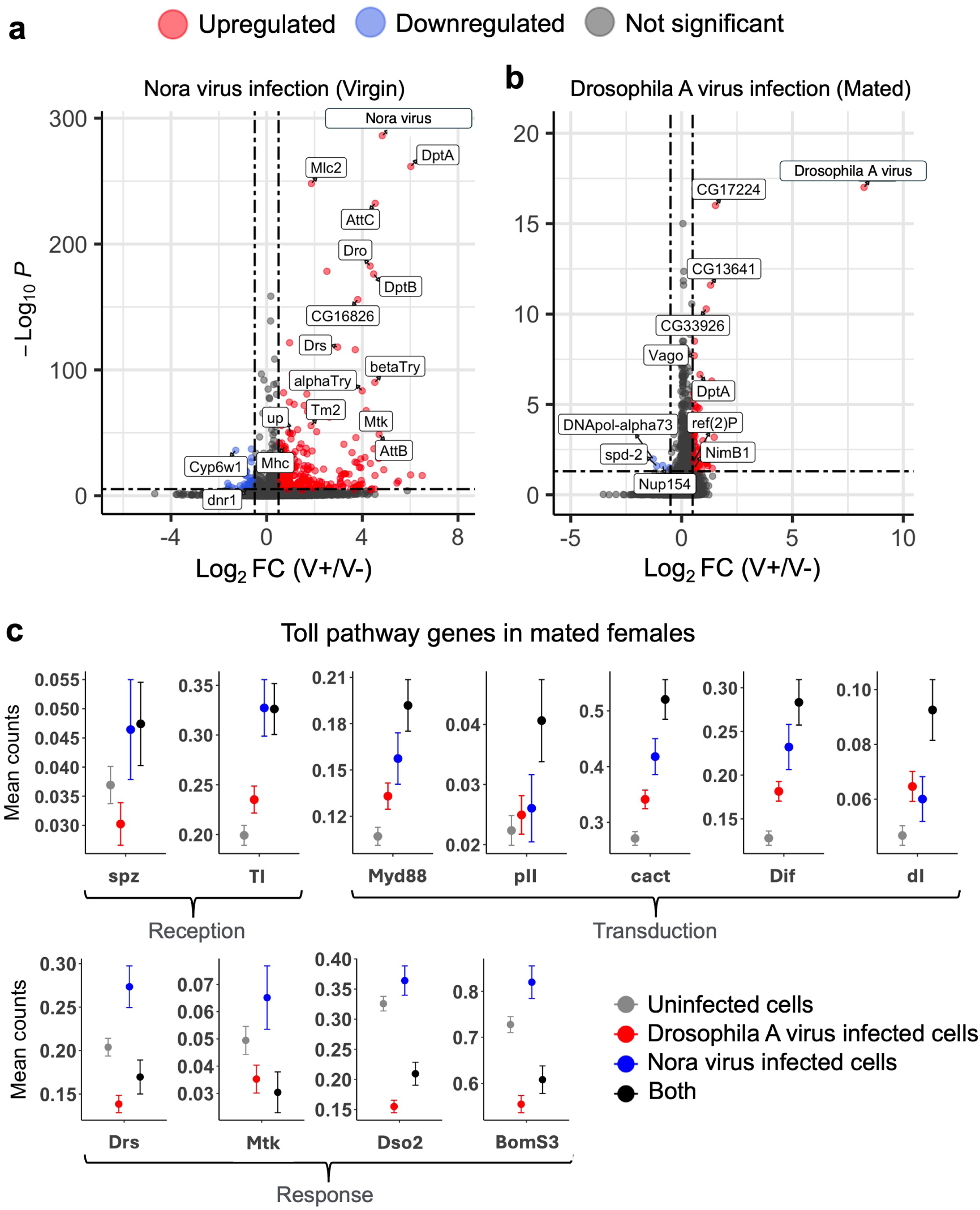
Differentially expressed gene (DEG) analysis during viral infection in the fat body. (a) Volcano plot of differentially expressed genes during Nora (virgin) infection. (0.05/total number of genes) was used as *pvalue* threshold. Absolute value of 0.5 was used as threshold for log_2_ fold change (Log_2_ FC). (b) Volcano plot of differentially expressed genes during Drosophila A virus (mated) infection. The same thresholds were applied as in panel (a). The analytical design for identifying differentially expressed genes, along with the statistical models applied, is summarized in Supplementary figure S29 and described in the methods section. (c) Mean counts of Toll pathway genes in Nora and Drosophila A virus infection in mated flies. Mean count was calculated from log normalized read counts from all the cells. Reads were normalized using Seurat’s LogNormalize method (counts per cell divided by total counts, scaled to 10,000, and log-transformed with log1p). The relation between log normalized values and CPM (counts per million) is (*e*^LogNormalized^ ^count^ − 1) × 100. Uninfected cells are those with zero RNA reads for both Nora and Drosophila A viruses. Drosophila A virus infected cells contain Drosophila A virus RNA reads (*>*0) but no Nora virus reads (0). Nora virus infected cells contain Nora virus RNA reads (*>*0) but no Drosophila A virus reads (0). Both means cells have both Nora and Drosophila A virus RNA reads(*>*0).

Enrichment analysis also revealed terms involving innate immunity and muscle contraction (Supplementary figure S13). In terms of immune response, we identified a total of 30 immune-related genes that were significantly differentially expressed in Nora virus infection (Supplementary figure S14, S15).

In strategy 2, we focused exclusively on replicate 2, the only virgin replicate with Nora virus infection, and compared uninfected cells to Nora virus-infected cells within only replicate 2. This analysis similarly revealed that IMD and Toll pathway genes were upregulated in association with Nora virus infection (Supplementary figure S16). We used strategy 1 for all the further analyses in this study for Nora virus infection in virgin flies.

In mated flies, replicate 1 had Nora virus infection, while replicate 2 had both Nora and Drosophila A virus. In this case, our strategy was to combine replicates 1 and 2. After combining, we compared virus uninfected cells with infected cells (see methods and Supplementary figure S29).

For Drosophila A virus infection in mated flies, we found that, the IMD pathway effector gene *DptA* and genes involved in antiviral immunity *(Vago, CG33926)* are upregulated while Toll pathway effector genes *(Drs, Mtk, Dso2, BomS3)* are downregulated (Figure 3b, c). The piRNA pathway suppresses TEs in germline cells, but recent studies have shown that in somatic cells like fatbody they are also expressed to suppress TEs [79]. Interestingly, the piRNA pathway gene *aub* was downregulated (−1.19, Log_2_ fold change, P-value=0.0033) in Drosophila A virus infection in mated flies. Supplementary figure S17 shows UMAP based visualization of upregulated and downregulated genes in Drosophila A virus infection. The gene ontology (GO) enriched terms identified cytoplasmic translation, ribosome assembly, ATP synthesis coupled proton transport (Supplementary figure S18). In terms of immune response, we identified a total of 8 immune-related genes that were significantly differentially expressed in Drosophila A virus infection (Supplementary figure S19).

One notable observation is that IMD and Toll pathway genes are upregulated in both virgin and mated flies in cells infected with Nora virus (Supplementary figures S20, S21). In contrast, in mated flies, the entire IMD pathway is upregulated in cells infected with Drosophila A virus (Supplementary figure S20). However, while the upstream components of the Toll pathway are upregulated, the effector genes (response), particularly Antimicrobial Peptides (AMPs) are downregulated in association with Drosophila A virus infection (Figure 3c). This suggests that Drosophila A virus may inhibit Toll effector gene transcription, perhaps by inhibition of *Dif* or *dl*. We also observed this pattern in cells that are coinfected with Nora and Drosophila A virus, meaning Drosophila A virus may regulate the impact of Nora virus (Figure 3c).

Additionally, in mated flies with Nora virus infection, we identified several differentially expressed genes, such as *Dredd* (antibacterial response), *eater* (phagocytosis), *Karl* (response to reactive oxygen species), *CYLD* (cell death), *Myo95E* (endocytosis) (Supplementary figures S22, S23). Cytoplasmic translation-related genes were found to be most enriched GO term (Supplementary figure S24). We identified 257 differentially expressed genes shared between Nora virus infected virgin and mated flies, including 13 immune-related genes (*Tsf1, Lectin-galC1, Fer2LCH, TotA, Thor, Dso1, Hml, pst, E(bx), SPE, Cdc42, NimB3, grh*). Among these shared genes, cytoplasmic translation genes such as *RpS23* and *RpL29* were most enriched. These genes are may be global response effect in Nora virus infection (Supplementary table ST8).

In conclusion, Nora and Drosophila A virus may activate canonical immune path-ways (IMD and Toll) in the fat body, and effector genes of the Toll pathway may be downregulated in association with Drosophila A virus infection, which could contribute to its higher viral titer in this tissue.

### 3.4 The response of each fat body cell type varies in magnitude during viral infection

Tissues consist of different cell types and each may be unique in terms of their function and response to pathogens. We wanted to understand cell specific gene expression to infection in the main three fat body cell types.

To study heterogeneity in response to Nora and Drosophila A virus infection, we examined how differentially expressed genes vary across cell types (interaction effect of cell type by virus), and used the interaction effect term of the ANCOVA model to formally assess cell type–by–infection effects for Nora virus in both virgin and mated flies, and for Drosophila A virus in the mated flies in the three identified fat body cell types.

In virgin flies (had only Nora virus infection), we found a total of 403 genes showing a cell cluster by virus infection interaction (Supplementary table ST5). Enrichment analysis revealed antibacterial humoral response, and defense response to gram positive bacterium are the two top enriched terms which also align with the enriched terms identified in the differentially expressed genes analysis (Supplementary figures S13 and S25). The response of top immune related genes that showed heterogeneity among fat body cell types is shown in Figure 4a. We observed broad upregulation of IMD pathway genes across nearly all cell types, although the magnitude of upregulation varied among cell types (Figure 4a). The log_2_ fold change of IMD pathway genes was higher in Fatbody cells 3, which is interestingly associated with the lower viral titer of Nora virus in these cells in virgin flies (Figure 2b). Notably, there were some genes that are up in some cells and down in others. For example, *CG6429* (humoral immune response) down in fat body cell 1 but up in fat body cell 2 and 3, *PGRP-LF* (IMD pathway gene) down in fat body cells 2, but up in fat body cells 1 and 3 (Figure 4a).

**Figure 4:**
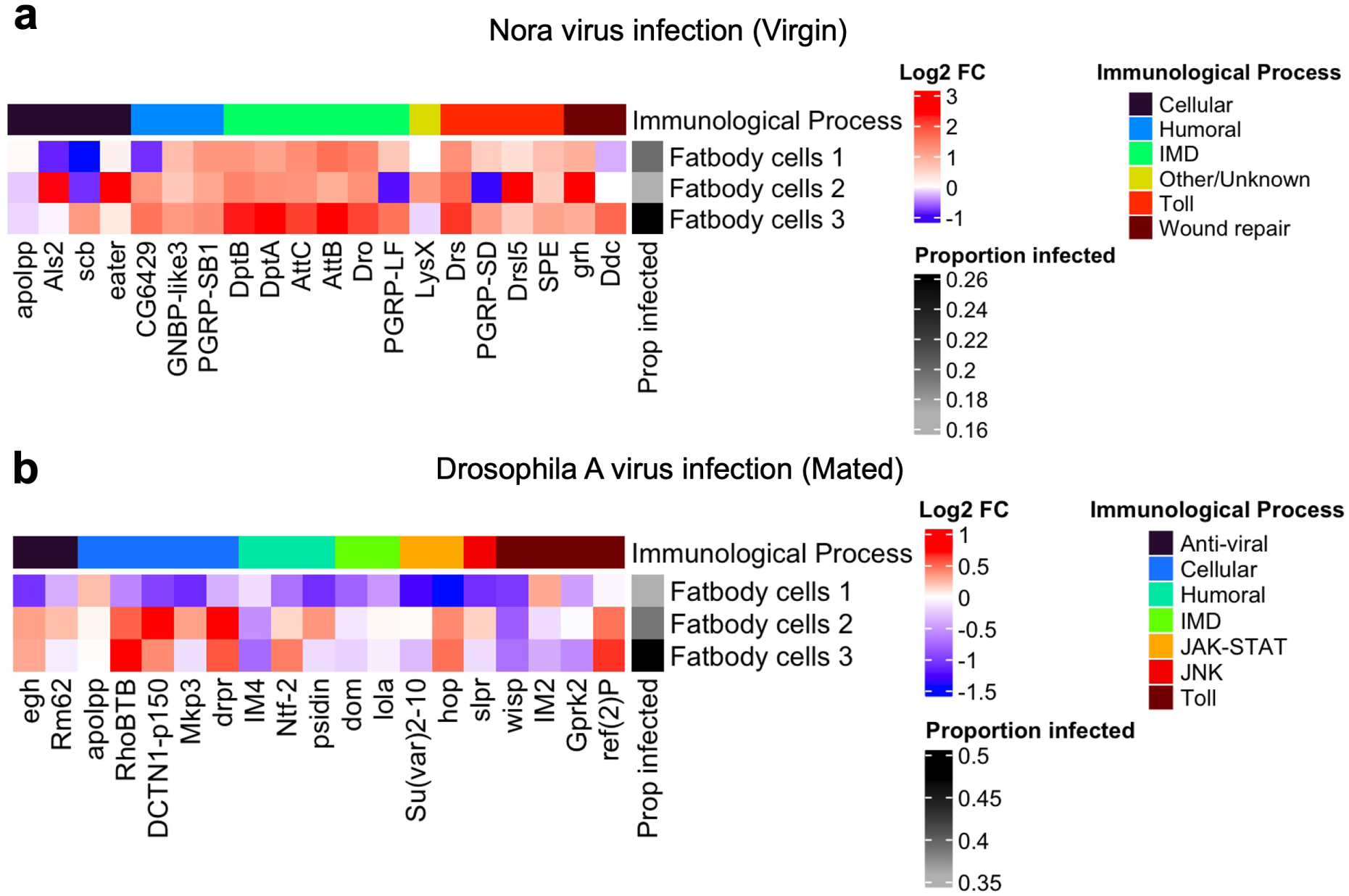
Cell type specific expression changes of immune related genes in viral infection in the fat body. (a) Log_2_ fold change (FC) of gene expression associated with Nora virus infection (infected/uninfected) in different cell types. Log_2_ fold change was calculated by comparing virus positive (viral RNA *>* 0) cells against virus negative cells (viral RNA=0). The percentage of virus-infected cells was determined by identifying cells with viral RNA read counts greater than zero. Top interaction effect immune genes (genes that change expression based cell type in infection) are shown. (b) Log_2_ fold change (FC) of gene expression associated with Drosophila A virus infection among different fat body cell types in mated flies. The same methods were applied to calculate Log_2_ fold change and the percentage of infected cells as in panel (a), and only the top immune related interaction effect genes are shown.

In mated flies, we found a total of 1578 genes showing a significant interaction effect between Drosophila A virus and cell type (Supplementary table ST5). Some of the genes were *dhd* (embryonic development), *Nplp2* (humoral immune response), *CG8745* (response to nicotine). Most of the top immune response genes with significant interaction effects exhibit heterogeneous expression across fat body cell types, with fat body cell type 1 showing predominantly downregulated immune responses, potentially explaining its higher Drosophila A virus titer (Figure 4b, 2b). Some genes showed opposite expression patterns across fat body cell types in Drosophila A virus infection as well. For example, *egh* was downregulated in fat body cell 1 but upregulated in fat body cells 2 and 3 (Figure 4b). Enrichment analysis of genes that had cell type by Drosophila A virus interaction effect, revealed nonassociative learning, and negative regulation of cellular macromolecule biosynthetic process are the two top enriched terms (Supplementary figure S26). In mated flies, we identified 39 genes (e.g., *UQCR-11L* - mitochondrial electron transport, *bgcn* - stem cell lineage differentiation) with significant interaction effects for Nora virus; however, this set did not yield any enriched Gene Ontology terms (Supplementary Table ST5).

In conclusion, viral infection elicits heterogeneous gene expression responses within the *Drosophila* fat body.

### 3.5 Dynamic regulation of transposable elements in viral infection

Recent studies have shown that in addition to germline activity, TEs are active in somatic cells [80, 81]. As noted earlier, piRNA pathway genes are produced in the *Drosophila* fat body and play a role in silencing TE activity within this tissue [79]. Additionally, Traffic jam (*tj*) is a master regulator of piRNA pathway in somatic cells [82]. *tj* in somatic cells can regulate the piRNA factor *flamenco*, thereby suppresses TE. In line with this, we observed that *flamenco* is upregulated in Nora virus infected cells in virgin flies (Supplementary figure S27) and piRNA pathway genes (*armi, aub*) are downregulated in Drosophila A virus infected cells of mated flies (Supplementary figure S17). RNAi pathway genes, which mediate antiviral immunity and suppress TEs in somatic cells [48], are also dysregulated during Nora virus and Drosophila A virus infection (Supplementary figure S28). We therefore examined whether TE transcript levels are altered in response to the viral infections.

We quantified DNA transposon, RC (Helitrons), LTR (retrotransposon) and LINE (retro-transposons). The expression of these TEs varied between virgin and mated flies in Nora virus infection (Figure 5a). In Nora virus infected cells, TE expression is downregulated in virgin flies but upregulated in mated flies. While piRNA pathway genes are known to protect against TEs in germline cells, they are also actively expressed in the *Drosophila* fat body (a somatic tissue) [79]. Given the different TE regulation patterns in virgin and mated flies, we examined the expression of two key piRNA pathway genes — one from the primary (*piwi, armi, vret, tj*) and one from the secondary (*AGO3, AGO1, aub, squ, vas*) pathway. We observed that in Nora virus infected cells of virgin flies, piRNA pathway genes are upregulated coincident with TE downregulation., whereas in mated flies they are predominantly downregulated, coinciding with TE upregulation (Figure 5b). The exact same pattern was also observed with RNAi pathway genes (Supplementary figure S28).

**Figure 5:**
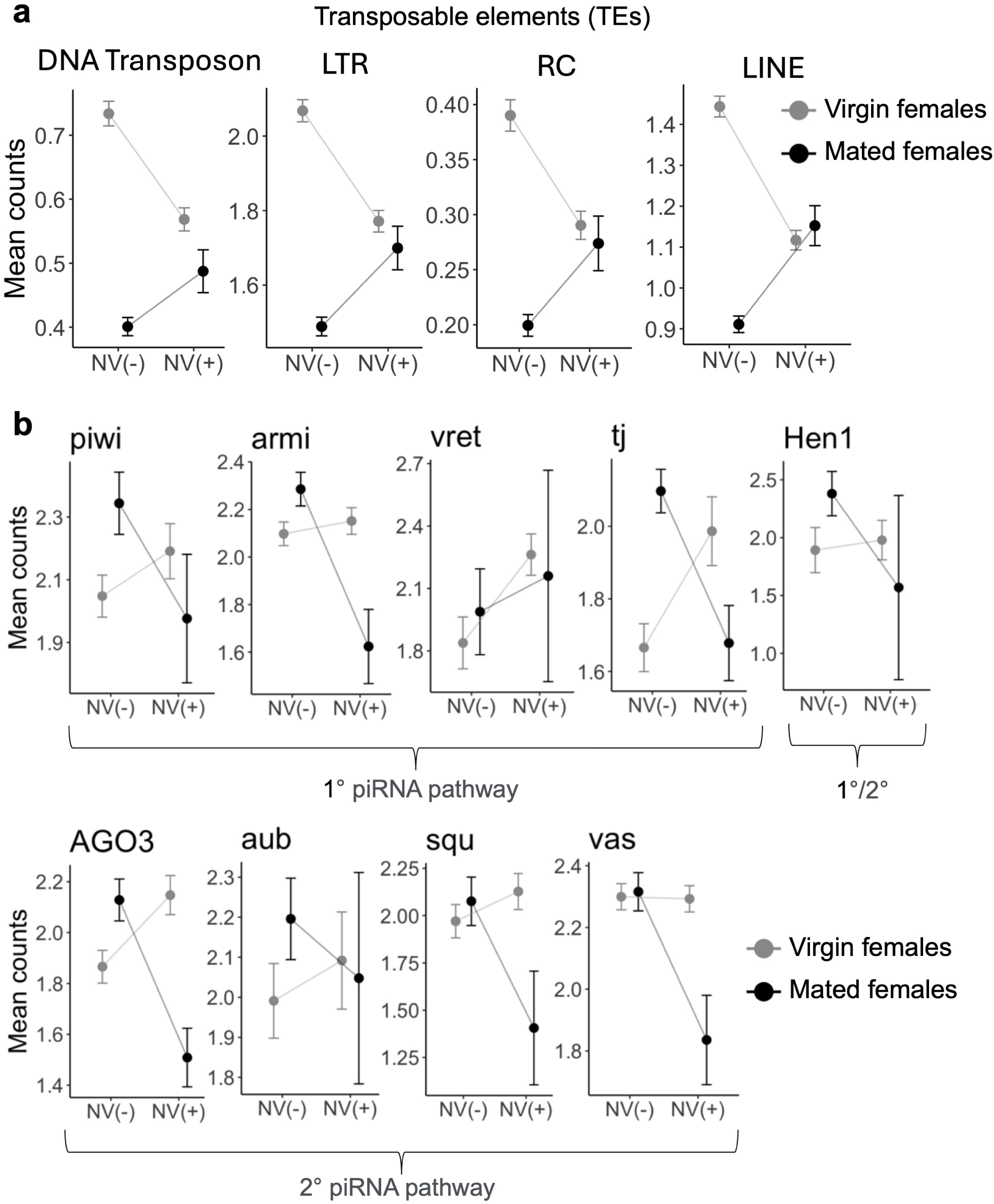
Transcriptional changes of TEs and piRNA pathway genes during Nora virus infection in the fat body. (a) Mean counts of TEs during Nora virus infection in virgin and mated flies. Mean counts are calculated by combining counts of all the cells. (b) Mean counts of piRNA pathway genes during Nora virus infection in virgin and mated flies. Mean counts are calculated by combining counts of the cells that expressed piRNA pathway genes. In general, counts are log normalized using Seurat’s LogNormalize method (counts per cell divided by total counts, scaled to 10,000, and log-transformed with log1p). NV(-) cells are those with no detectable Nora virus RNA (viral RNA reads = 0). NV(+) cells are those with Nora virus RNA reads *>* 0 and no Drosophila A virus RNA (reads = 0), specifically in the mated flies where both viruses are present.

Somatic TE expression in virgin versus mated flies has been rarely investigated. We observed that in uninfected cells, TE expression is lower in mated flies compared to that of virgins (Figure 5a) and interestingly, under conditions of Nora virus infection, mated flies show a reversal of this downregulation. This may reflect a loss of host control over TEs or stress release TEs during viral infection (Figure 5a).

Conversely, TE expression is upregulated in Drosophila A virus infection and the piRNA pathway genes are downregulated (Figure 6a,b). However, tissues from virgin females were not infected with Drosophila A virus, so we cannot examine the effect of mating. Overall, it is difficult to reconcile why TE expression increases with viral infection in mated flies but decreases in virgins, and how the piRNA and RNAi pathways contribute to this pattern. Prior study showed that piRNA pathway is not required for antiviral defense [83]. So, we hypothesize that, TE dysregulation is likely due to RNAi pathway as it’s dual role in antiviral immunity and suppressing TEs in somatic cells. When the RNAi pathway protects against viruses by regulating its core components *(AGO2, Dcr-2)*, as an artefact TEs are also affected.

**Figure 6:**
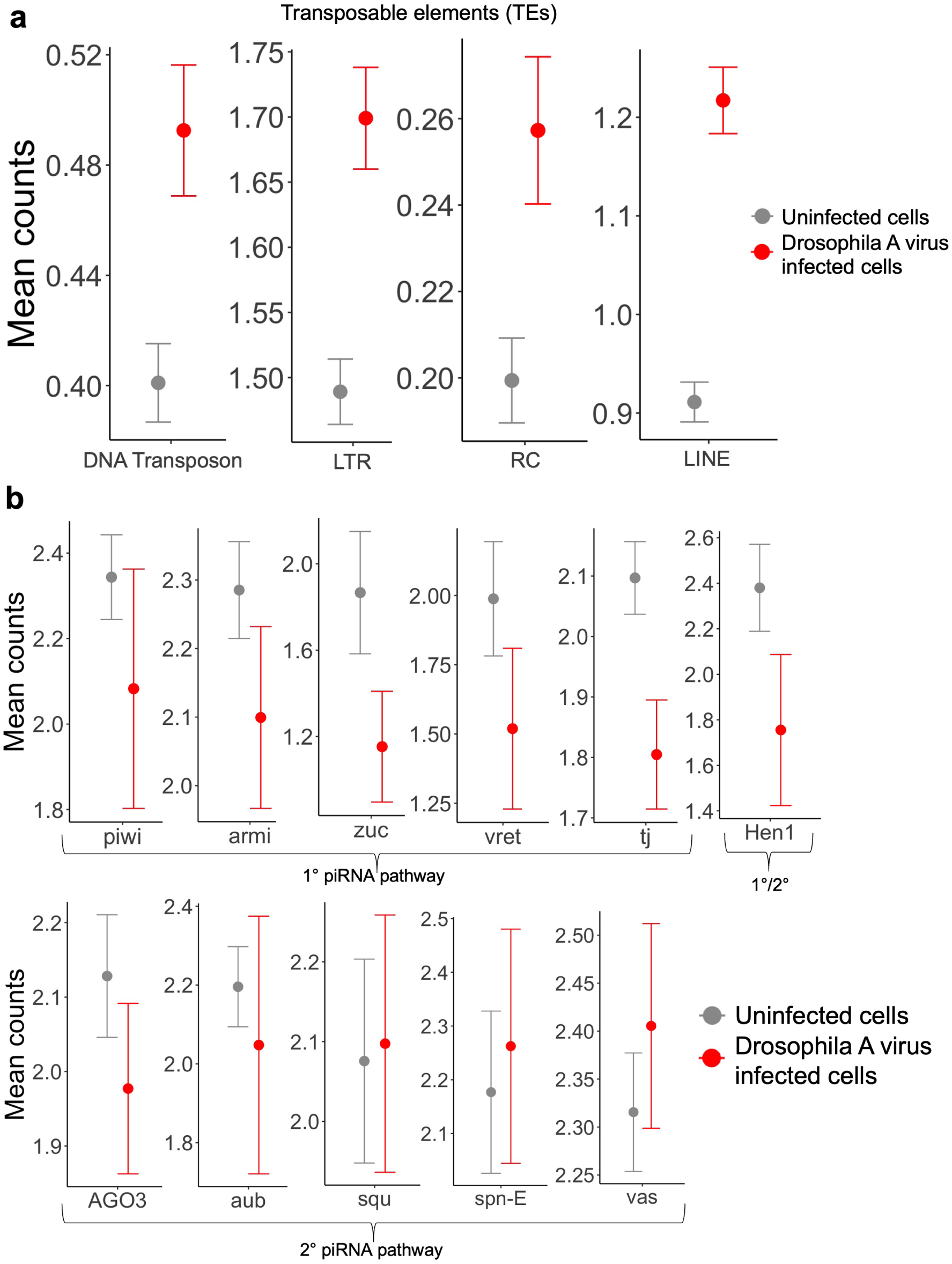
Transcriptional changes of TEs and piRNA pathway genes associated with Drosophila A virus infection in the fat body. (a) Mean counts of TEs associated with Drosophila A virus infection in mated flies. Mean counts are calculated by combining counts of all the cells. (b) Mean counts of piRNA pathway genes during Drosophila A virus infection in mated flies. Mean counts are calculated by combining counts of the cells that expressed piRNA pathway genes. Counts are log normalized for both panel a and b. Seurat’s LogNormalize method was used (counts per cell divided by total counts, scaled to 10,000, and log-transformed with log1p). Uninfected cells are those with no detectable Drosophila A virus RNA (viral RNA reads = 0). Drosophila A virus infected cells are those with Drosophila A virus RNA reads *>* 0 and no Nora virus RNA (reads = 0), as in mated flies both viruses are present.

In conclusion, viral infection may influence TE transcript levels in the somatic fat body of *Drosophila*, potentially reflecting a loss of host regulation of TEs.

## 4 Discussion

We studied cell tropism, differential gene responses, and TE dynamics of Nora virus and Drosophila A virus infection in the *Drosophila* fat body using publicly available single-cell RNA sequencing data. Both viruses infect a variety of fat body cell types; however, Drosophila A virus exhibits a higher viral titer and infects a greater number of cells compared to Nora virus during coinfection in mated flies. This disparity suggests that Drosophila A virus may have evolved strategies to more effectively evade or suppress host defenses, leading to enhanced viral proliferation. Previous studies have implicated the Toll pathway in host responses to both Drosophila A virus and Nora virus [29, 32]. Here our study indicate that, unlike Nora virus, Drosophila A virus may suppress Toll pathway effector genes (AMPs), a mechanism that could explain this phenomenon (Figure 3c). This may also suggest that during coinfection, the *Drosophila* fat body is more effective at suppressing Nora virus, which creates an opportunity for Drosophila A virus to spread efficiently. A recent study showed that Drosophila A virus infects multiple Drosophilidae host species at higher viral titers than Nora virus, a finding that complements our observations [84]. Interestingly, we observed a negative correlation between viral titer and the number of infected cells among cell types (Supplementary figure S12). Among the three major fat body cell types, Fatbody 3 exhibited the highest number of infected cells but the lowest viral titer for both viruses, regardless of condition (Figure 2). This pattern may be explained by the trade-off hypothesis of pathogen virulence, which posits that excessively high viral titers can limit cell-to-cell transmission by inducing host cell death, whereas maintaining lower, optimal titers may enhance transmission potential [85, 86]. A similar phenomenon has been reported in a study of influenza virus [87].

Several studies have mentioned that TE transcript levels even in somatic cells can be altered during infection caused by different pathogens such as viruses [39, 41]. Consistent with this, we observed differences in TE dynamics between mated and virgin flies during persistent viral infection(Figure 5,6). In virgin flies infected with Nora virus, TE expression is generally downregulated, driven by the upregulation of the piRNA pathway, which is essential for silencing TEs (Figure 5a,b). However, in mated flies infected with Nora virus, the piRNA pathway is downregulated, resulting in increased TE expression which may indicate that Nora virus infection can restore TE expression to pre-mating levels. For Drosophila A virus infection in mated flies, we observed TE upregulation. (Figure 6).

Previous studies show that these viruses start their infection primarily in the gut but as the gut barrier gets disrupted, it allows these viruses to spread into other tissues as well [28, 88]. Further studies in additional *Drosophila* tissues are needed to clarify viral tropism across different tissue types.

## 5 Conclusion

We characterized the transcriptional landscape associated with persistent viral infection in the *Drosophila* fat body at single nucleus level. Our results indicate that persistent viruses are broadly distributed across fat body cell types, although susceptibility varies among specific populations. Host transcriptional responses to viral infection are not restricted to a subset of cell types; rather, transcriptional responses are shared across all fat body cell types, with the magnitude of activation differing between cell types. Additionally, alterations in TE activity suggest that persistent viral infection is associated with changes in genome regulatory dynamics within somatic tissues. Overall, our study provides insights into viral cell tropism, gene, and TE expression pattern in persistent viral infections in the *Drosophila* fat body.

## Acknowledgements

We thank Justin P. Blumenstiel and members of the Unckless lab for valuable feedback.

## Author information

**Department of Molecular Biosciences, The University of Kansas, USA.**

Nilanjan Roy, Robert L. Unckless

## Contributions

Conceptualization: NR, RLU; Data curation: NR, RLU; Formal analysis: NR, RLU; Funding acquisition: RLU; Investigation: NR, RLU; Methodology: NR, RLU; Project administration: NR, RLU; Visualization: NR, RLU; Writing: NR, RLU.

## Corresponding authors

Correspondence to Nilanjan Roy or Robert L. Unckless.

## Supplementary information

**Additional file, SM1:** Supplementary figures.

**Additional table, ST1:** Marker genes of the *Drosophila* fat body cell types.

**Additional table, ST2:** Gene ontology (GO) analysis of the marker genes of *Drosophila* fat body cell types.

**Additional table, ST3:** Probability of infection of the fat body cell types.

**Additional table, ST4:** Nora and Drosophila A virus expression comparison in different fat body cell types.

**Additional table, ST5:** Interaction effect genes in the fat body cells in viral infection.

**Additional table, ST6:** Differential gene expression analysis in Nora virus infection in virgin flies.

**Additional table, ST7:** Differential gene expression analysis in Nora and Drosophila A virus infection in mated flies.

**Additional table, ST8:** Differentially expressed genes shared between virgin and mated flies in Nora virus infection.

## Availability of data and material

The processed data and code will be made available in the peer-reviewed publication.

