## Supplementary materials for "Persistent viral infection in the *Drosophila* fat body is associated with immune activation at the single cell level"

### 1 Additional figures

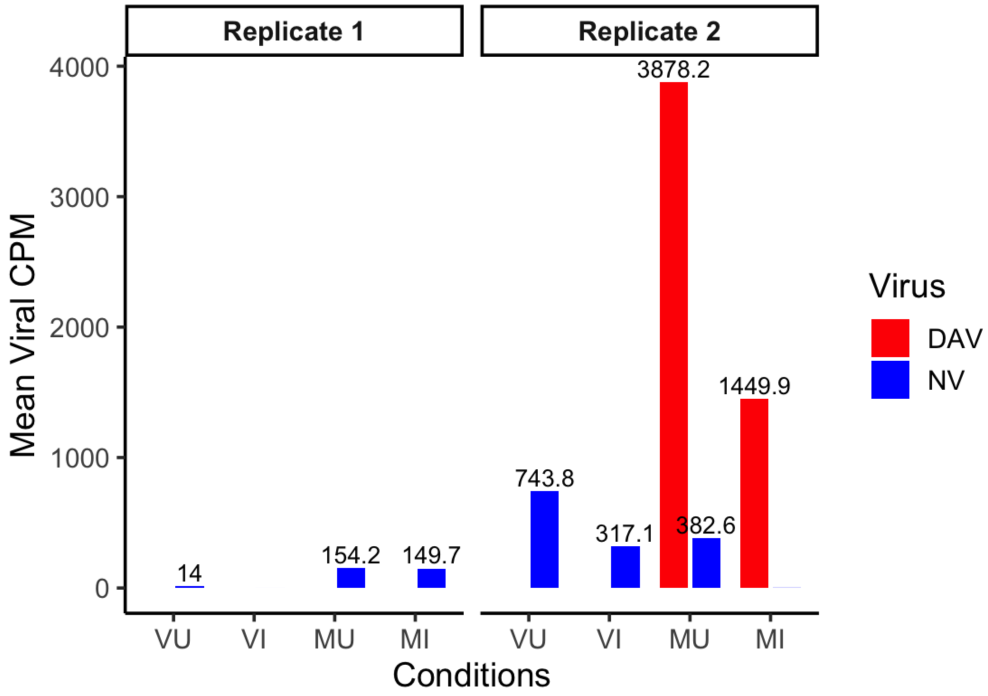


Figure S1: **Mean viral RNA counts across all experimental conditions.** Replicate 1 of the MU and MI groups had substantial Nora virus (NV) RNA counts. Drosophila A virus (DAV) RNA was detected only in Replicate 2 (MU and MI). Nora virus reads in Replicate 2 were observed in VU, VI, and MU samples. CPM denotes counts per million. Abbreviations: VU = Virgin bacterial uninfected; VI = Virgin bacterial infected; MU = Mated bacterial uninfected; MI = Mated bacterial infected. VU, MU was used for all the subsequent analysis. Refer to the methods section for details regarding the replicates. DAV = Drosophila A virus; NV = Nora virus.

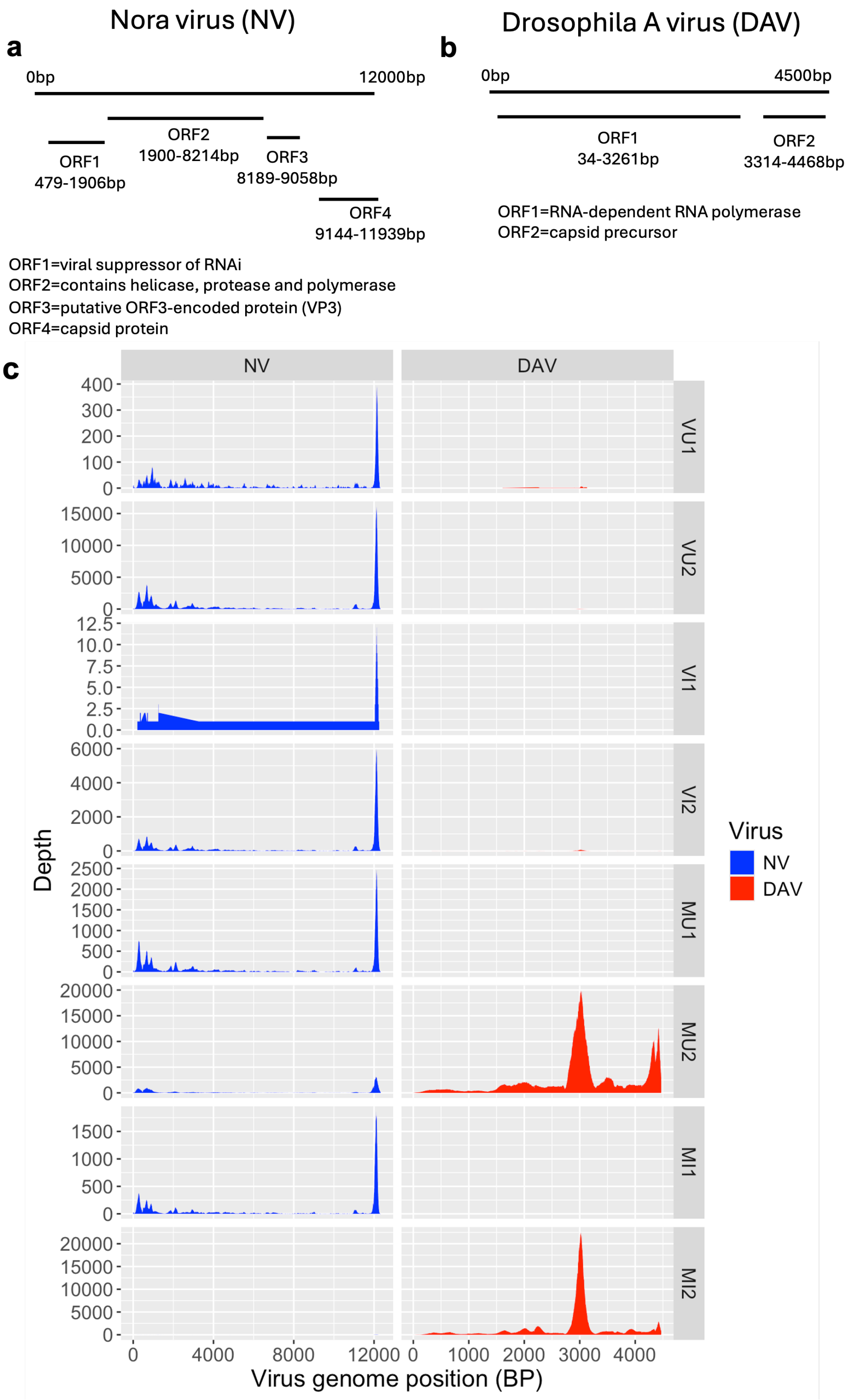


Figure S2: **Depth at each Nora and Drosophila A virus genomic position across different conditions and replicates.** (c) Depth indicates the number of reads. Reads were derived using samtools depth command. The majority of the Nora virus reads were aligned to ORF1 (viral suppressor of RNAi pathway) and ORF4 (capsid protein). Similarly, most reads mapped to ORF2 (capsid protein) of Drosophila A virus. Abbreviations: NV=Nora virus, VU=Virgin bacterial uninfected, VI=Virgin bacterial infected, MU=Mated bacterial uninfected, MI=Mated bacterial infected. VU, MU was used for all the subsequent analysis. Refer to the methods section for details regarding the replicates. BP= Base pair. (a), (b) represents the genomes of NV and DAV.

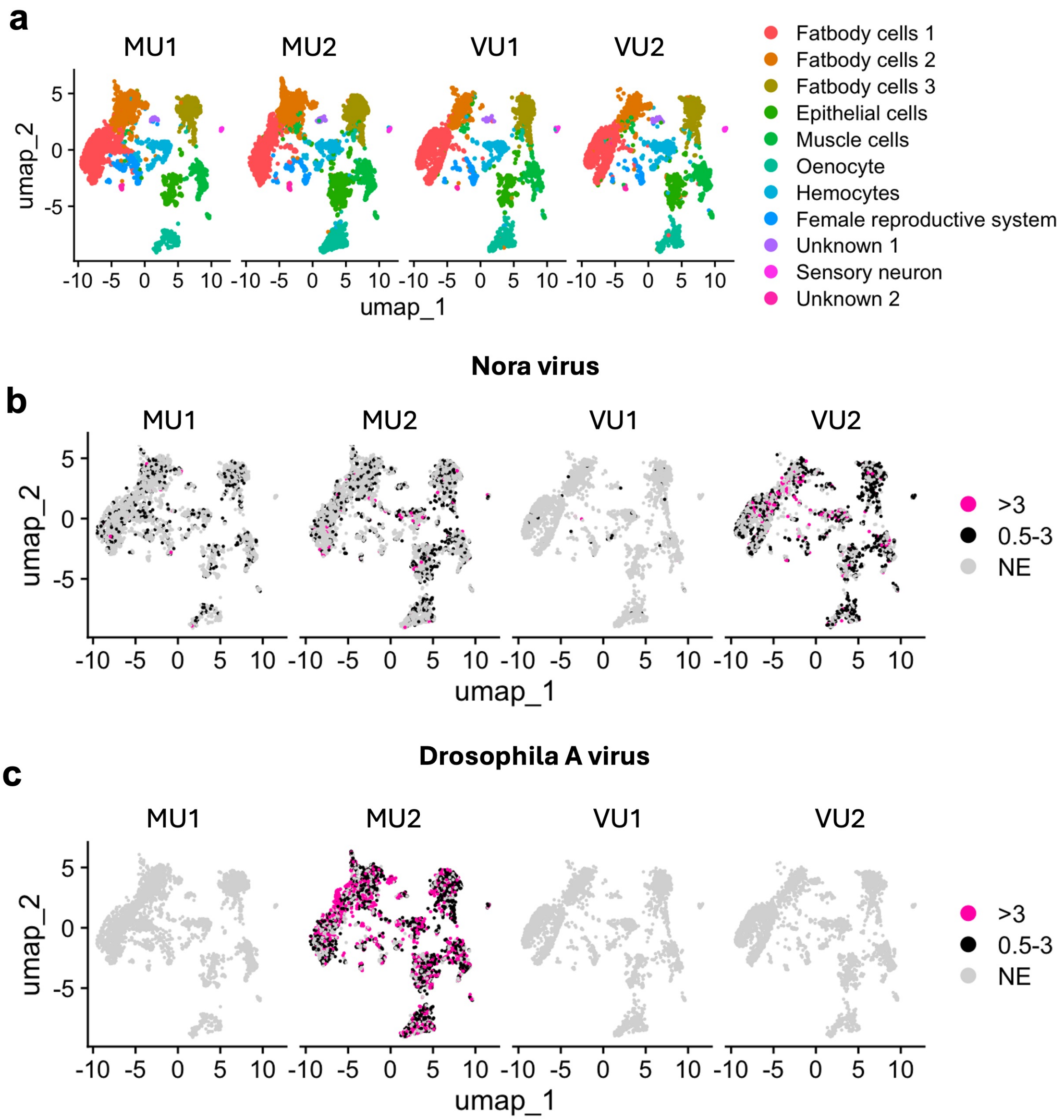


Figure S3: **Nora and Drosophila A virus tropism in fat body cell types.** (a) UMAP plot of 11 fat body-associated clusters in the replicates of virgin and mated flies. Colors and numbers in the UMAP indicate respective inferred cell types. VU=Virgin bacterial uninfected, VI=Virgin bacterial infected, MU=Mated bacterial uninfected, MI=Mated bacterial infected. VU, MU was used for all the subsequent analysis. Refer to the methods section for details regarding the replicates. (b) Nora virus tropism in different fat body cell types in mated and virgin flies. (c) Drosophila A virus tropism in different fat body cell types in mated and virgin flies. Read counts are log normalized. Seurat LogNormalize method was used where feature counts for each cell are divided by the total counts of that cell, multiplied by a scale factor of 10,000, and then taking natural log–transformed value using log1p. >3 represents cells where viral read counts are greater than 3. 0.5-3 represents cells where viral read counts are between 0.5 to 3. NE (not expressed) represents cells that don't have any viral RNA. Log normalized counts can be converted to CPM using the formula $cpm = (e^{\text{LogNormalized count}} - 1) \times 100$. Thus, values $> 3$ correspond to viral RNA reads greater than $\sim$2000 CPM, while values between 0.5 and 3 correspond to viral RNA reads between $\sim$65–2000 CPM.

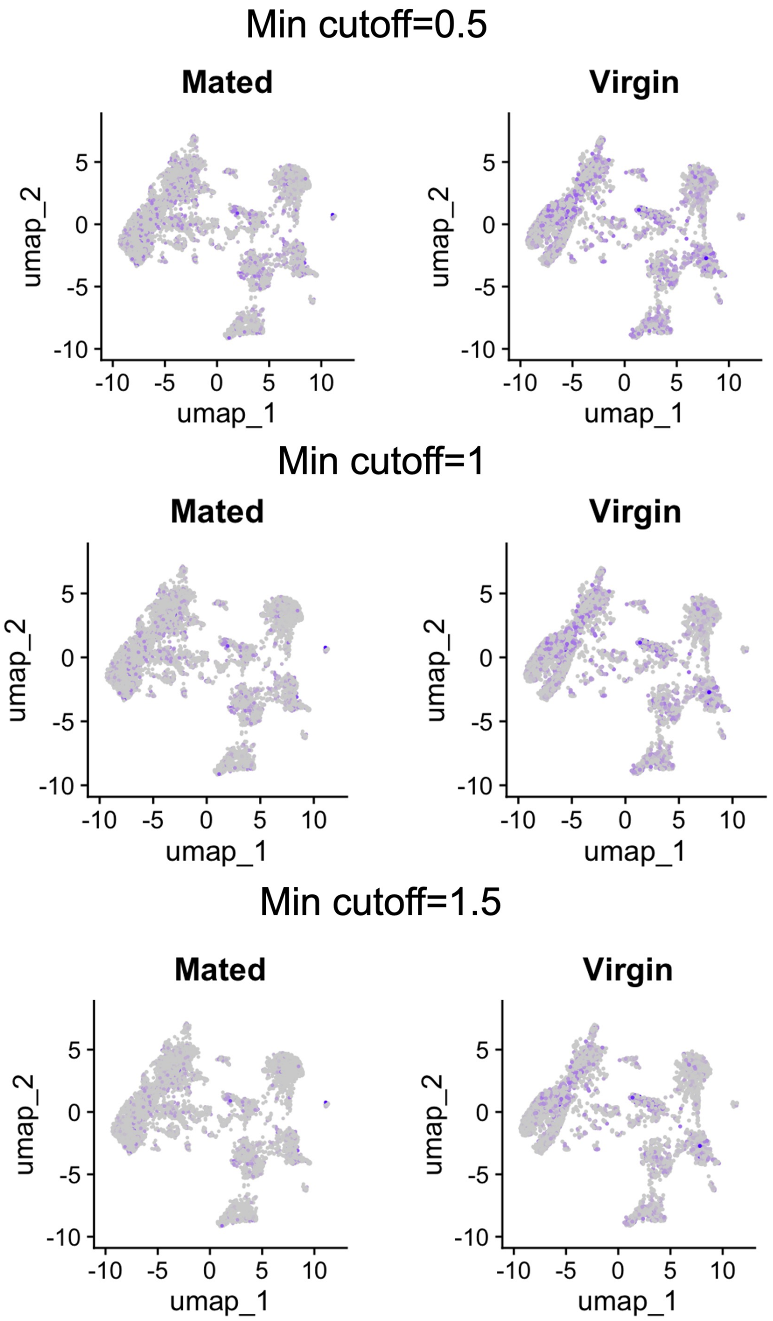


Figure S4: **Nora virus tropism across various fat body cell types at different viral titer thresholds.** Viral RNA read counts was used as proxy for viral titer. Colored cells have viral titer of atleast the minimum cutoff. Nora virus RNA is distributed across different cell types and it is consistent at different viral titer cutoffs. Read counts are log normalized. Seurat LogNormalize method was used where feature counts for each cell are divided by the total counts of that cell, multiplied by a scale factor of 10,000, and then taking natural log–transformed value using log1p. Minimum cutoff 0.5 represents cells where viral read counts are at least 0.5. Minimum cutoff 1 represents cells where viral read counts are at least 1. Minimum cutoff 1.5 represents cells where viral read counts are at least 1.5. Log normalized counts can be converted to CPM (counts per million) using the formula $CPM = (e^{\text{LogNormalized count}} - 1) \times 100$. Thus, 0.5, 1, 1,5 correspond to viral RNA reads of 65, 172, 348 CPM respectively.

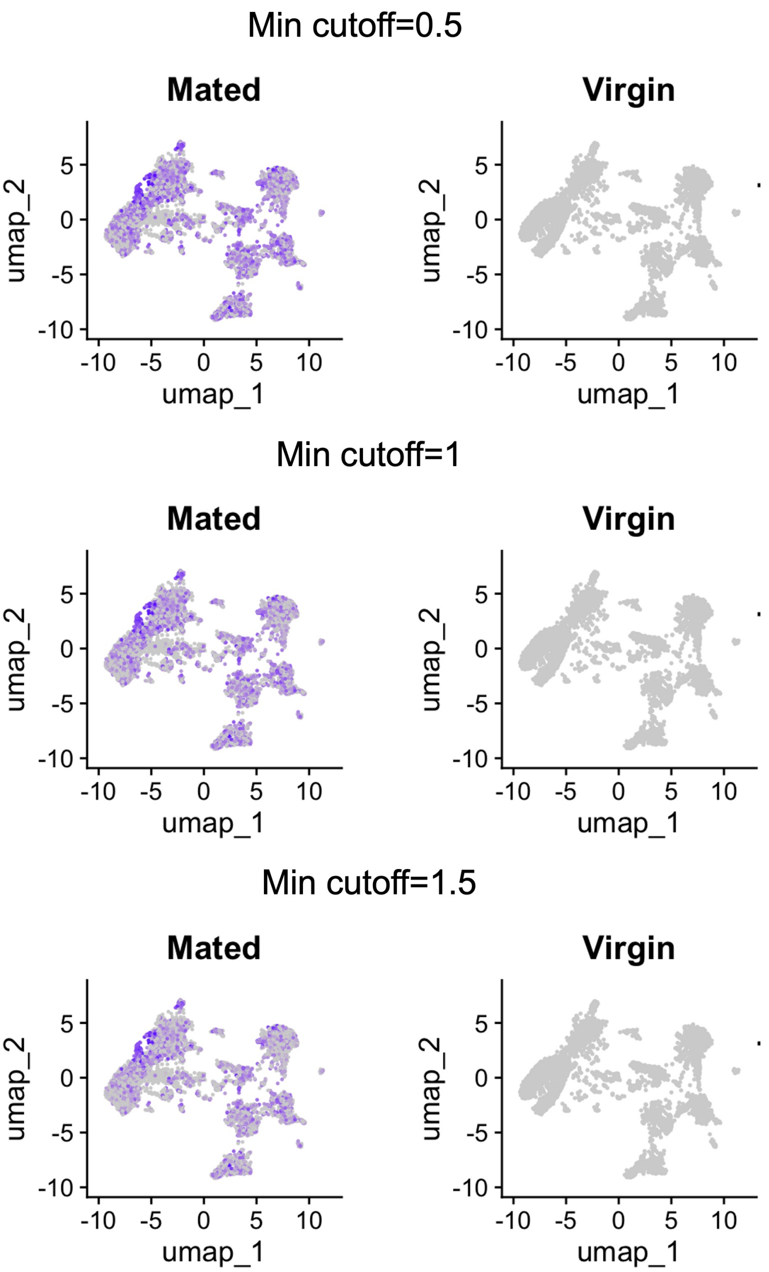


Figure S5: **Drosophila A virus tropism in different cell types of fat body at different viral titer cutoff.** Viral RNA read counts was used as proxy for viral titer. Colored cells have viral titer of atleast the minimum cutoff. Similarly like Nora virus, Drosophila A virus RNA is distributed across different cell types and it is consistent at different viral titer cutoffs as well. Read counts are log normalized. Seurat LogNormalize method was used where feature counts for each cell are divided by the total counts of that cell, multiplied by a scale factor of 10,000, and then taking natural log–transformed value using log1p. Minimum cutoff 0.5 represents cells where viral read counts are at least 0.5. Minimum cutoff 1 represents cells where viral read counts are at least 1. Minimum cutoff 1.5 represents cells where viral read counts are at least 1.5. Log normalized counts can be converted to CPM (counts per million) using the formula $CPM = (e^{\text{LogNormalized count}} - 1) \times 100$. Thus, 0.5, 1, 1,5 correspond to viral RNA reads of 65, 172, 348 CPM respectively.

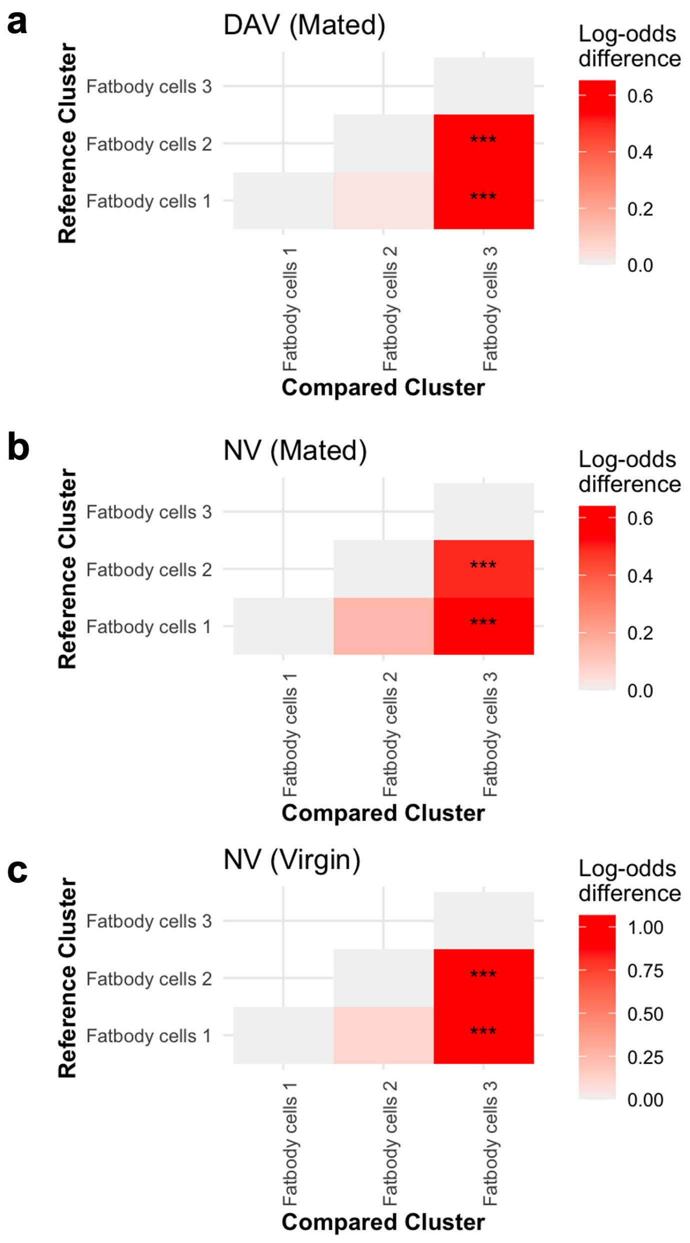


Figure S6: **Infection probability of fat body cell types**. (a) Infection probability of fat body cell types in Drosophila A virus infection in mated flies. GLM model: (DAV infected cells, DAV uninfected cells) $\sim$ cell types+NV infection+Replicates. (b) Infection probability of fat body cell types Nora virus infection in mated flies. GLM model: (NV infected cells, NV uninfected cells) $\sim$ cell types+DAV infection+Replicates. (c) Infection probability of fat body cell types Nora virus infection in virgin flies. GLM model: (NV infected cells, NV uninfected cells) $\sim$ cell types+DAV infection+Replicates. The model were fitted using a binomial family with a logit link function and followed by post hoc tukey test to compare infection probabilty between cell types. Redness (higher log odd difference) means that the compared cluster/cell type has higher infection odds than the reference cluster/cell type. Asterisks indicates statistically significant. Uninfected cells are those with zero RNA reads for both Nora and Drosophila A viruses. DAV infected cells contain Drosophila A virus RNA reads (>0) but no Nora virus reads (0). NV infected cells contain Nora virus RNA reads (>0) but no Drosophila A virus reads (0). DAV= Drosophila A virus, NV= Nora virus.

Model (glm): NV(0/1) ~ DAV(0/1) + Replicates + Cell types
0=Virus uninfected cells, 1=virus infected cells

| | Estimate | Std. Error | z value | Pr(>\|z\|) |
|---|---|---|---|---|
| (Intercept) | -1.79856 | 0.06103 | -29.472 | < 2e-16 *** |
| DAV(0/1) | 0.39936 | 0.07514 | 5.315 | 1.07e-07 *** |

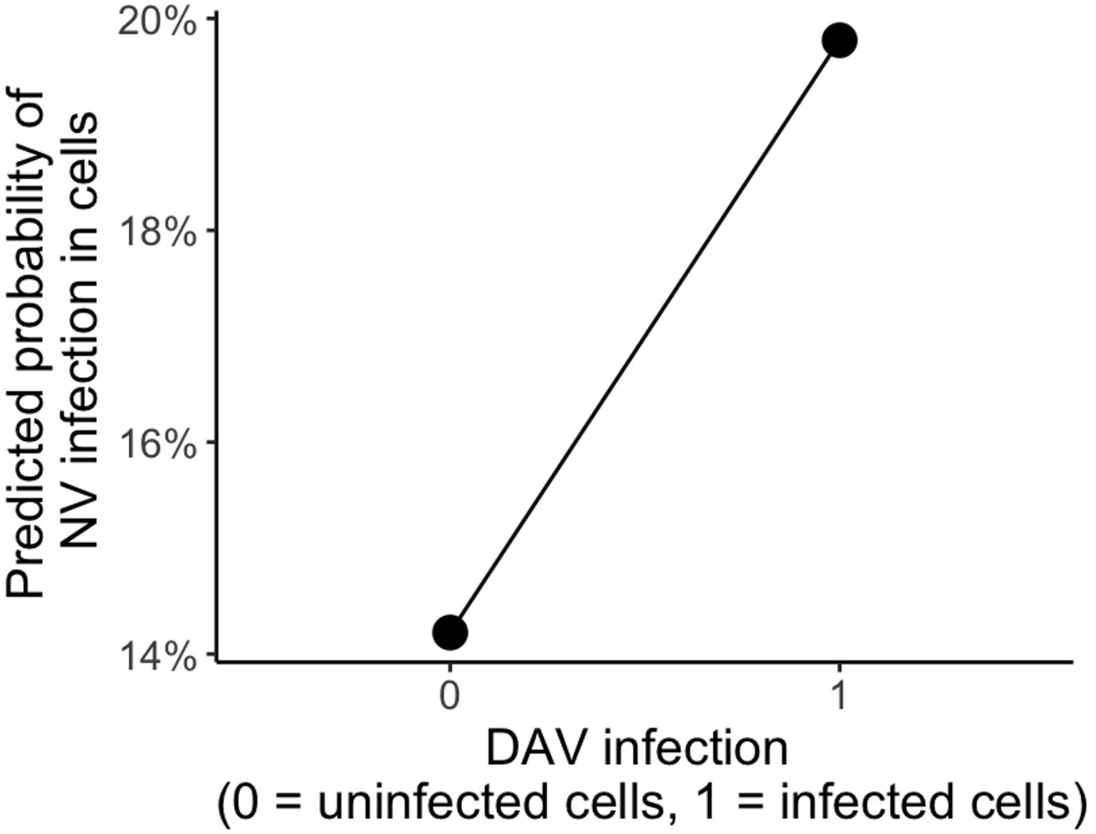


Figure S7: **Coinfection probabilty of Nora and Drosophila A virus in fat body cells**. GLM model: NV(0/1) $\sim$ DAV(0/1) + Replicates + Cell types. If a cell is infected with Drosophila A virus then the cell has higher chance of having Nora virus. Uninfected cells are those with zero RNA reads for both Nora and Drosophila A viruses. DAV infected cells contain Drosophila A virus RNA reads (>0) but no Nora virus reads (0). NV infected cells contain Nora virus RNA reads (>0) but no Drosophila A virus reads (0). DAV= Drosophila A virus, NV= Nora virus.

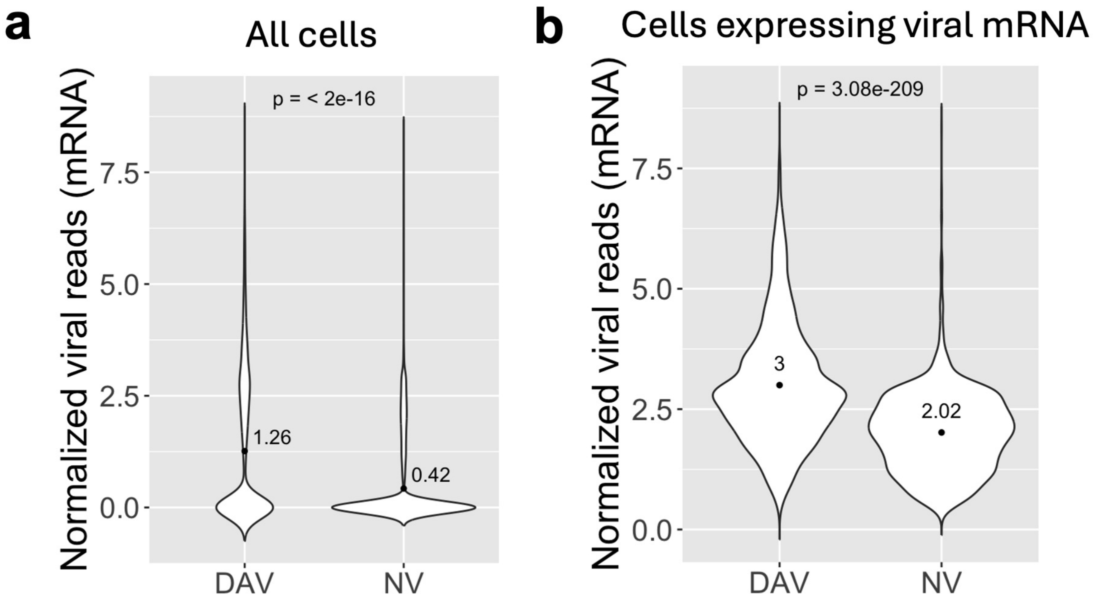


Figure S8: **Comparison of Nora and Drosophila A virus normalized reads (viral mRNA) in whole fat body tissue in mated flies**. (a) Nora and Drosophila A virus normalized reads across all the fat body cells (whole tissue). (b) Nora and Drosophila A virus normalized reads in cells expressing the virus, viral RNA reads >0 (whole tissue). In both cases, Drosophila A virus reads are significantly higher than the Nora virus. Wilcoxon test was used to compare Nora and Drosophila A virus reads. Read counts are log normalized. Seurat LogNormalize method was used where feature counts for each cell are divided by the total counts of that cell, multiplied by a scale factor of 10,000, and then taking natural log–transformed value using log1p. For CPM (counts per million) reference, log normalized counts can be converted to CPM using the formula $CPM = (e^{\text{LogNormalized count}} - 1) \times 100$. DAV= Drosophila A virus, NV= Nora virus.

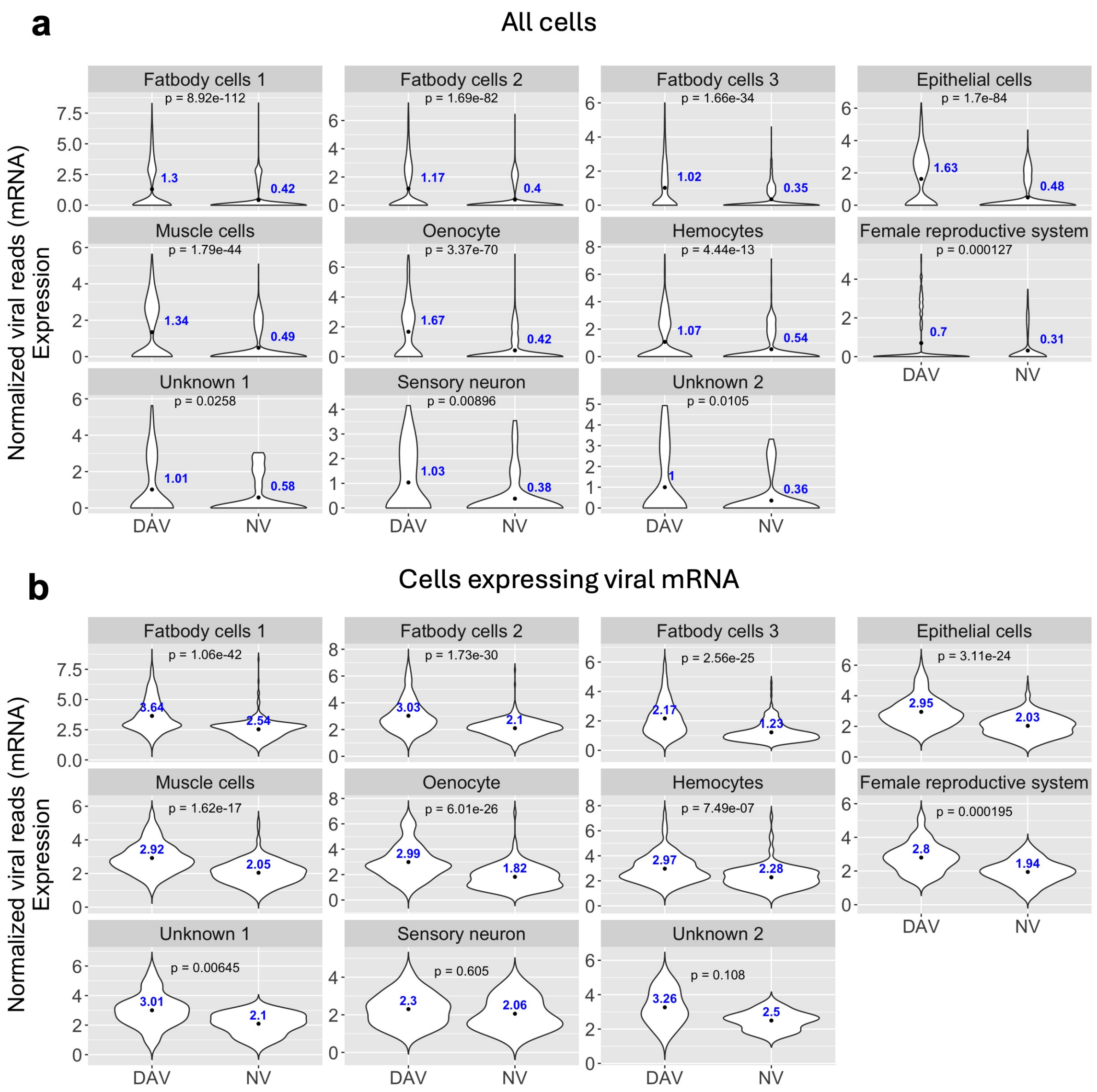


Figure S9: **Comparison of Nora and Drosophila A virus normalized reads (viral RNA) in fat body cell types in mated flies**. (a) Nora and Drosophila A virus normalized reads across all the fat body cell types. (b) Nora virus and Drosophila A virus normalized reads across all the fat body cell types where the viral RNA is expressed (viral RNA reads >0). In both cases, the majority of the cell types have higher Drosophila A virus reads than Nora virus. The ANOVA framework (posthoc tukey) was used to compare Nora and Drosophila A virus reads. Read counts are log normalized. Seurat LogNormalize method was used where feature counts for each cell are divided by the total counts of that cell, multiplied by a scale factor of 10,000, and then taking natural log–transformed value using log1p. For CPM (counts per million) reference, log normalized counts can be converted to CPM using the formula $CPM = (e^{\text{LogNormalized count}} - 1) \times 100$. DAV= Drosophila A virus, NV= Nora virus.

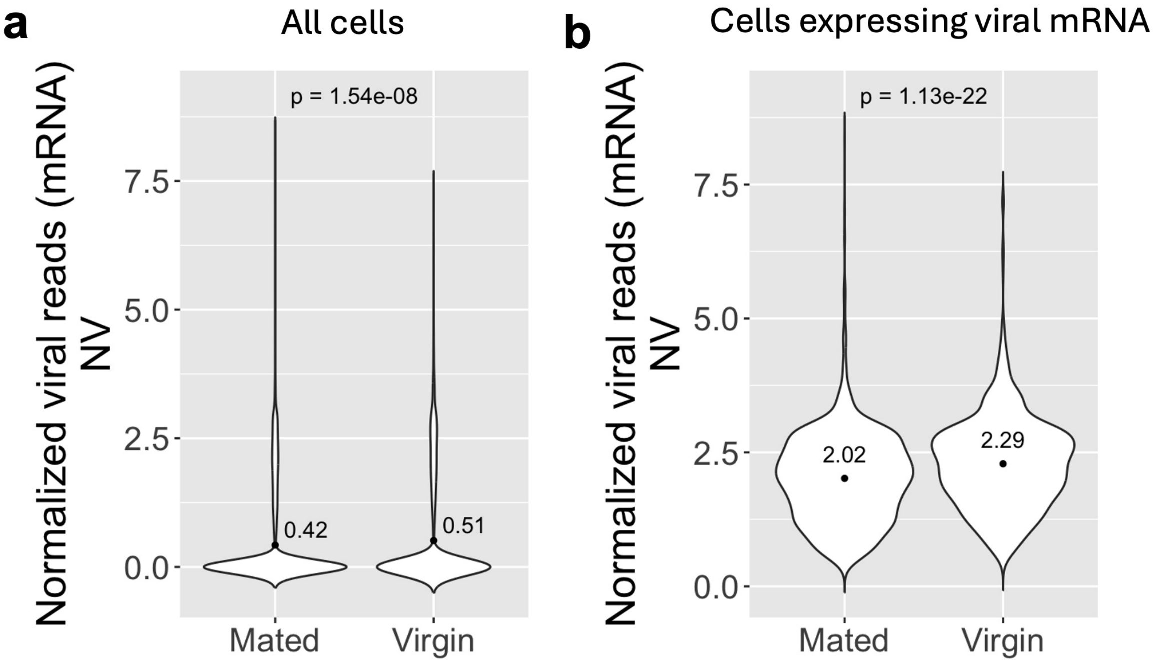


Figure S10: **Comparison of Nora virus normalized reads (viral RNA) in fat body tissue between mated and virgin flies**. (a) Nora virus normalized reads across all the fat body cells in mated and virgin flies. (b) Nora virus normalized reads in cells expressing the virus (viral reads >0) in mated and virgin flies. For both (a), and (b) virgin flies had higher Nora virus reads. Wilcoxon test was used to compare Nora virus reads between mated and virgin condiitons. Read counts are log normalized. Seurat LogNormalize method was used where feature counts for each cell are divided by the total counts of that cell, multiplied by a scale factor of 10,000, and then taking natural log–transformed value using log1p. For CPM (counts per million) reference, log normalized counts can be converted to CPM using the formula $CPM = (e^{\text{LogNormalized count}} - 1) \times 100$. NV=Nora virus.

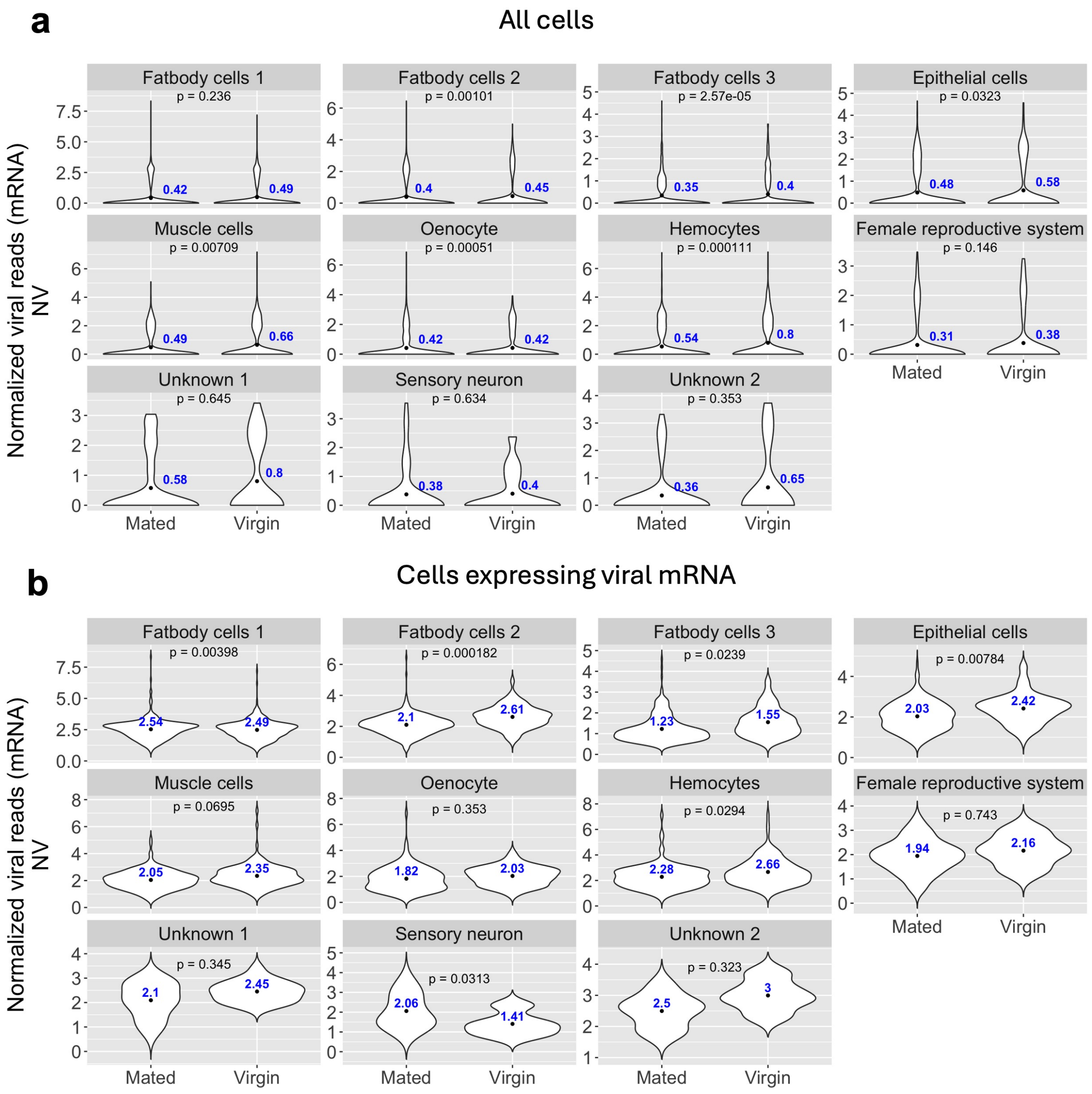


Figure S11: **Comparison of Nora virus normalized reads (viral RNA) in fat body cell types between mated and virgin flies.**(a) Nora virus normalized reads across different fat body cell types. (b) Nora virus normalized reads in different fat body cell types where the viral RNA is expressed (viral RNA reads >0). For both (a), and (b) virgin flies had higher Nora virus reads in majority of the cell types. The ANOVA framework (posthoc tukey) was used to compare Nora virus reads between mated and virgin flies. Read counts are log normalized. Seurat LogNormalize method was used where feature counts for each cell are divided by the total counts of that cell, multiplied by a scale factor of 10,000, and then taking natural log–transformed value using log1p. For CPM (counts per million) reference, log normalized counts can be converted to CPM using the formula $CPM = (e^{\text{LogNormalized count}} - 1) \times 100$. NV=Nora virus.NV=Nora virus.

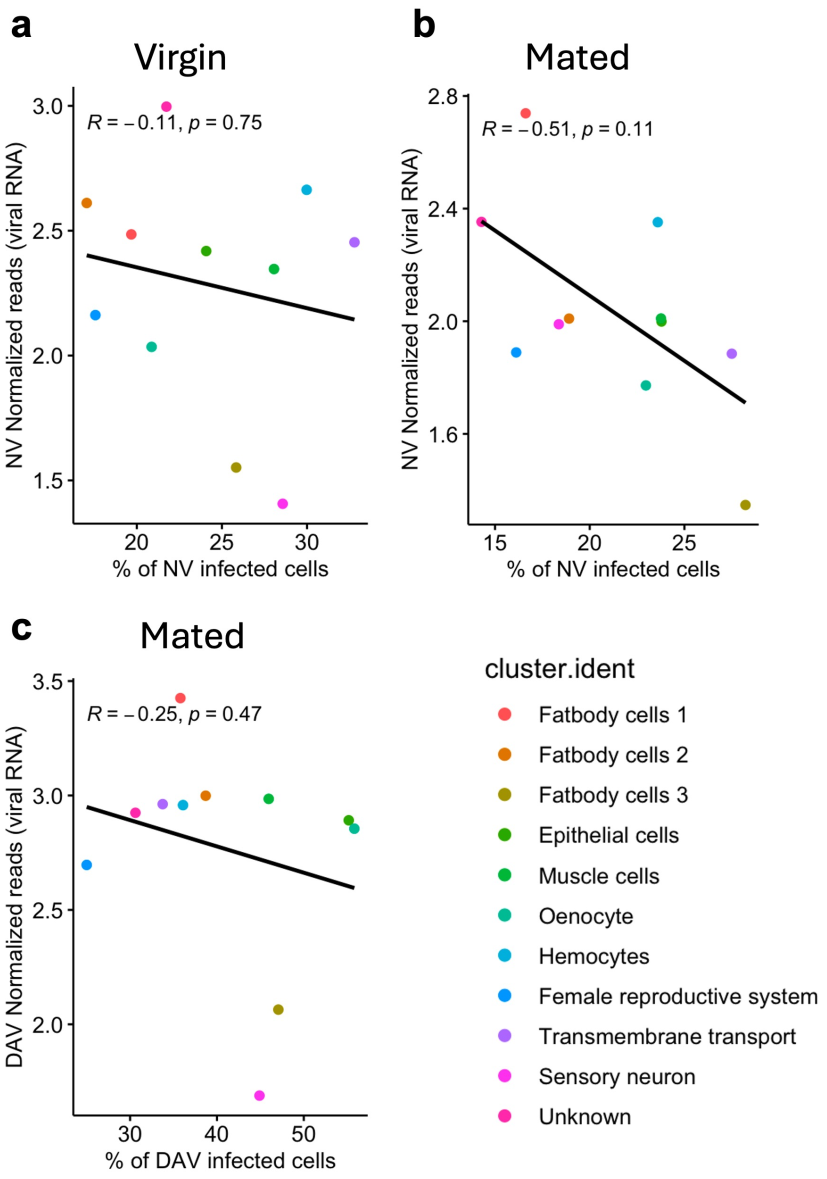


Figure S12: **Relationship between the percentage of infected cells with viral titer in the fat body.** (a) Relationship between the percentage of cells infected with Nora virus and viral titer in fat body cell types in virgin flies. (b) Relationship between the percentage of cells infected with Nora virus and viral titer in fat body cell types in mated flies. (c) Relationship between the percentage of cells infected with Drosophila A virus and viral titer in fat body cell types in mated flies. Although not statistically significant, in all cases we found a negative correlation between viral titer and percentage of infected cells. Normalized viral reads were used as a proxy for viral titer. The percentage of infected cells was determined by considering cells as infected when viral reads were $> 0$. Read counts are log normalized using Seurat LogNormalize method, where feature counts for each cell are divided by the total counts of that cell, multiplied by a scale factor of 10,000, and then taking natural log–transformed value using log1p. For CPM (counts per million) reference, log normalized counts can be converted to CPM using the formula $CPM = (e^{\text{LogNormalized count}} - 1) \times 100$. DAV= Drosophila A virus, NV= Nora virus.

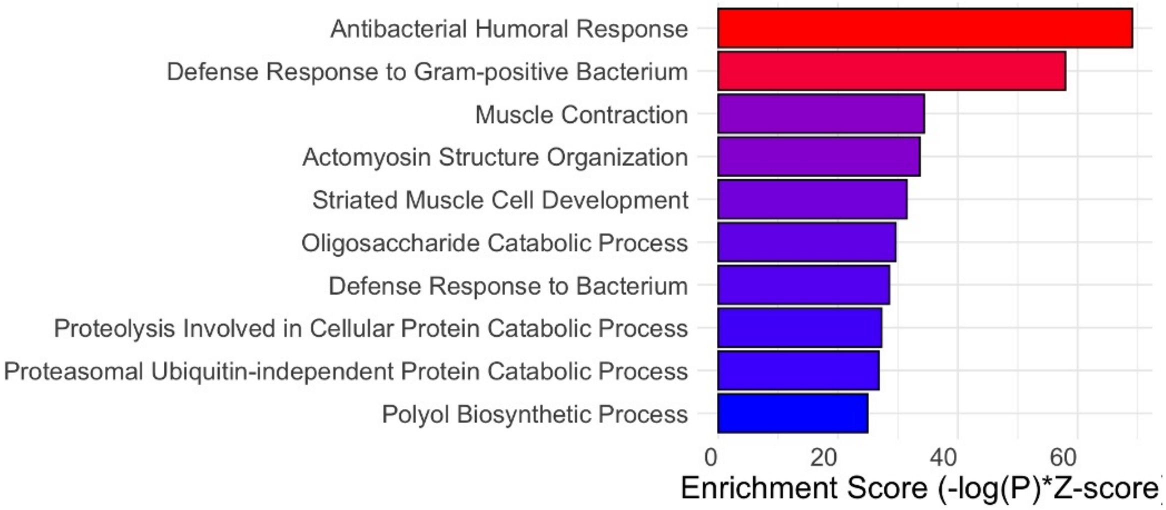


Figure S13: **Gene ontology (GO) enrichment analysis of differentially expressed genes in virgin flies in Nora virus infection.** Differentially expressed gene set was selected using a p-value threshold of $1 \times 10^{-5}$ for the GO analysis.

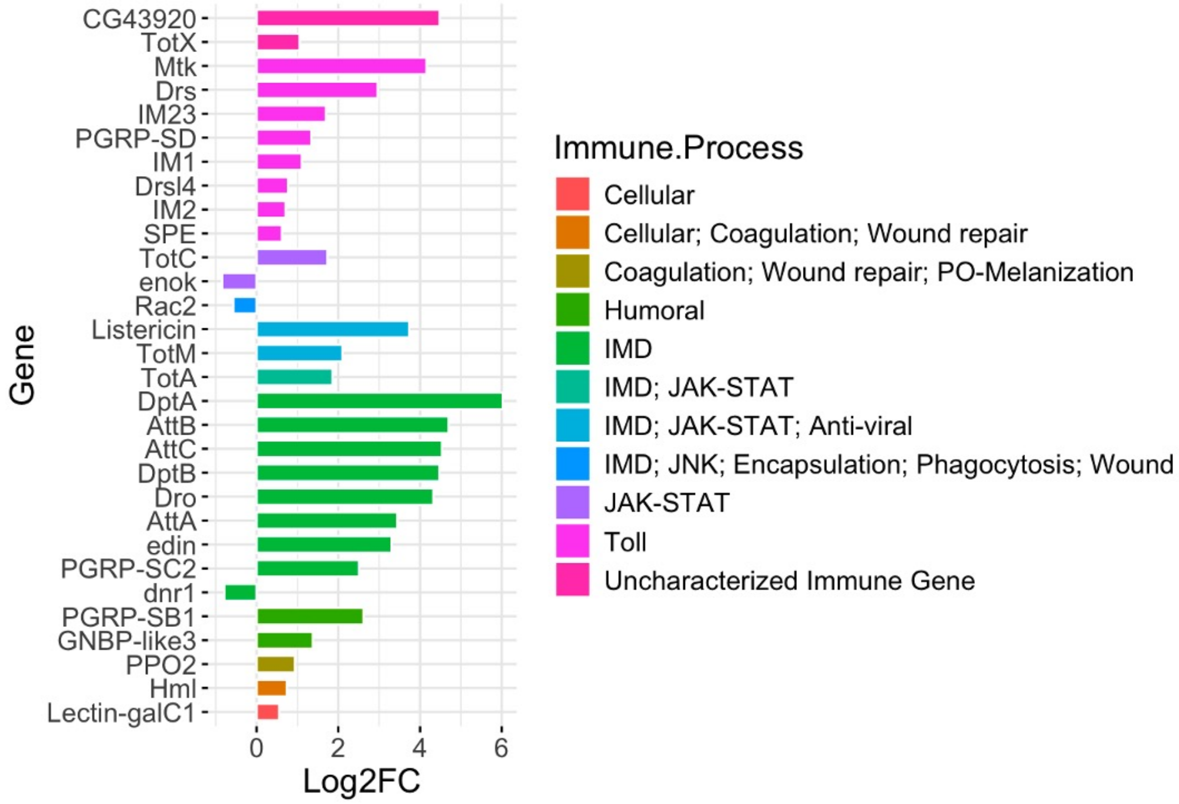


Figure S14: **Differentially expressed immune genes in virgin flies in Nora virus infection.** The majority of the immune genes are upregulated in association with Nora virus infection. The gene set was selected using a p-adjusted (BH method) threshold of 0.05 and an absolute $\text{Log}_2$ fold change cutoff of 0.5. $\text{Log}_2\text{FC} = \text{Log}_2$ fold change.

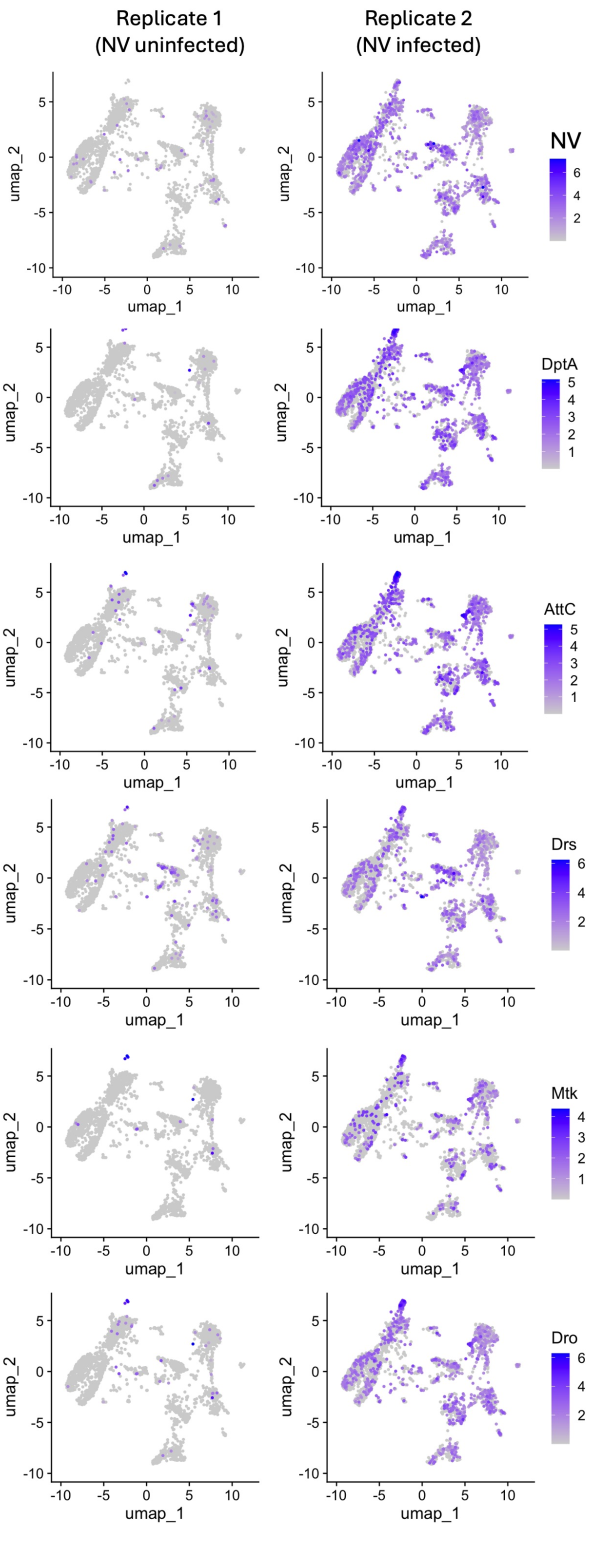


Figure S15: **UMAP-based comparison of differentially expressed genes between uninfected and Nora virus infected flies (virgin) in the fat body.** Immune related genes are upregulated. Read counts are log normalized using Seurat LogNormalize method, where feature counts for each cell are divided by the total counts of that cell, multiplied by a scale factor of 10,000, and then taking natural log–transformed value using log1p. For CPM (counts per million) reference, log normalized counts can be converted to CPM using the formula $CPM = (e^{\text{LogNormalized count}} - 1) \times 100$. NV= Nora virus.

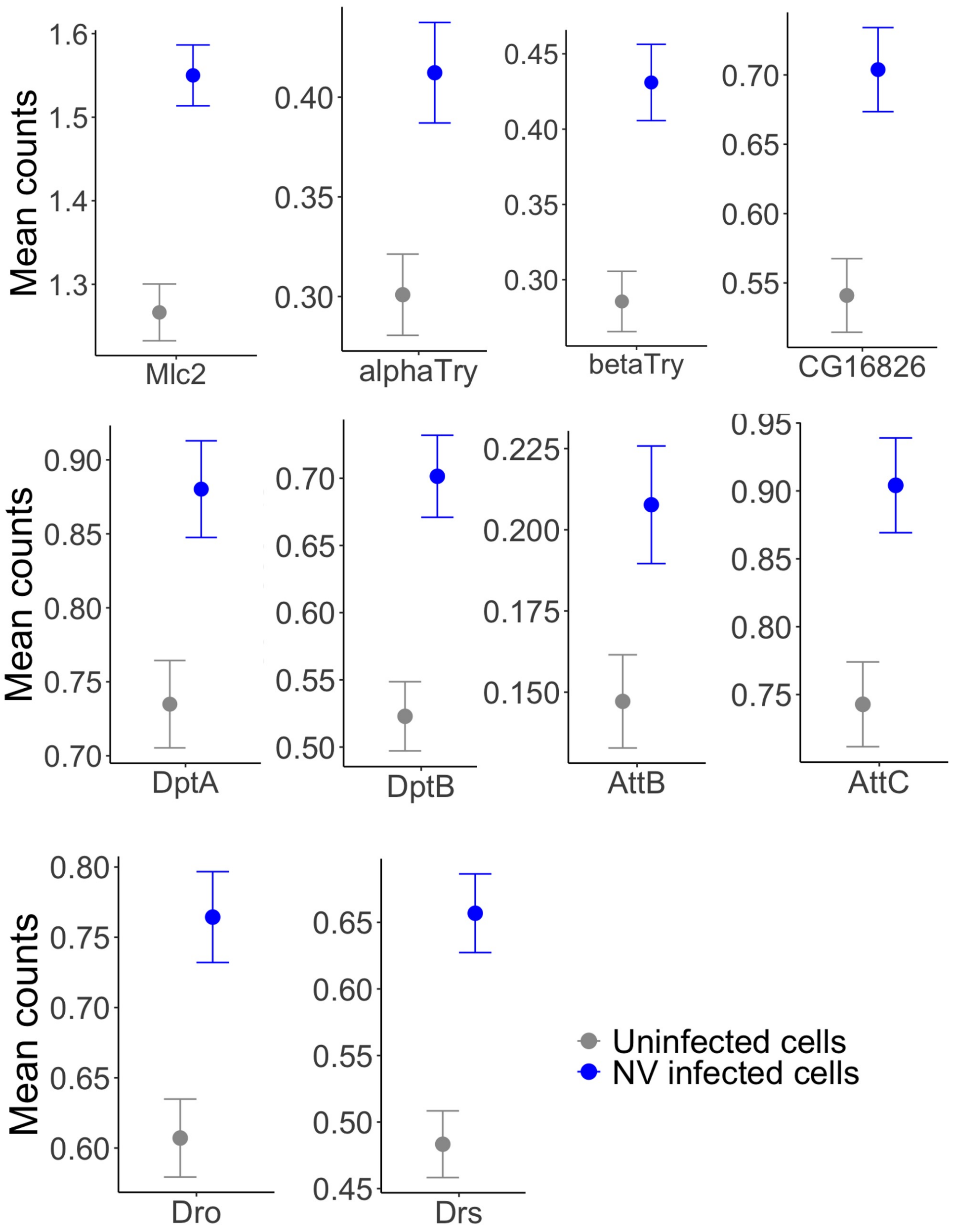


Figure S16: **Mean counts of differentially expressed genes (identified using method 1) in Nora virus uninfected versus infected cells, analyzed exclusively in Nora virus infected virgin flies (replicate 2).** This represents method 2. In this approach, the genes identified as upregulated in method 1 were also found to be upregulated, demonstrating colinearity between the two methods. This similarity increases confidence in the identified differentially regulated genes. Refer to the results section for further details on both methods. Uninfected cells are those with no detectable Nora virus RNA (viral RNA reads = 0). Nora infected cells are those with Nora virus RNA reads > 0. Read counts are log normalized using Seurat LogNormalize method, where feature counts for each cell are divided by the total counts of that cell, multiplied by a scale factor of 10,000, and then taking natural log–transformed value using log1p. For CPM (counts per million) reference, log normalized counts can be converted to CPM using the formula $CPM = (e^{\text{LogNormalized count}} - 1) \times 100$. NV= Nora virus.

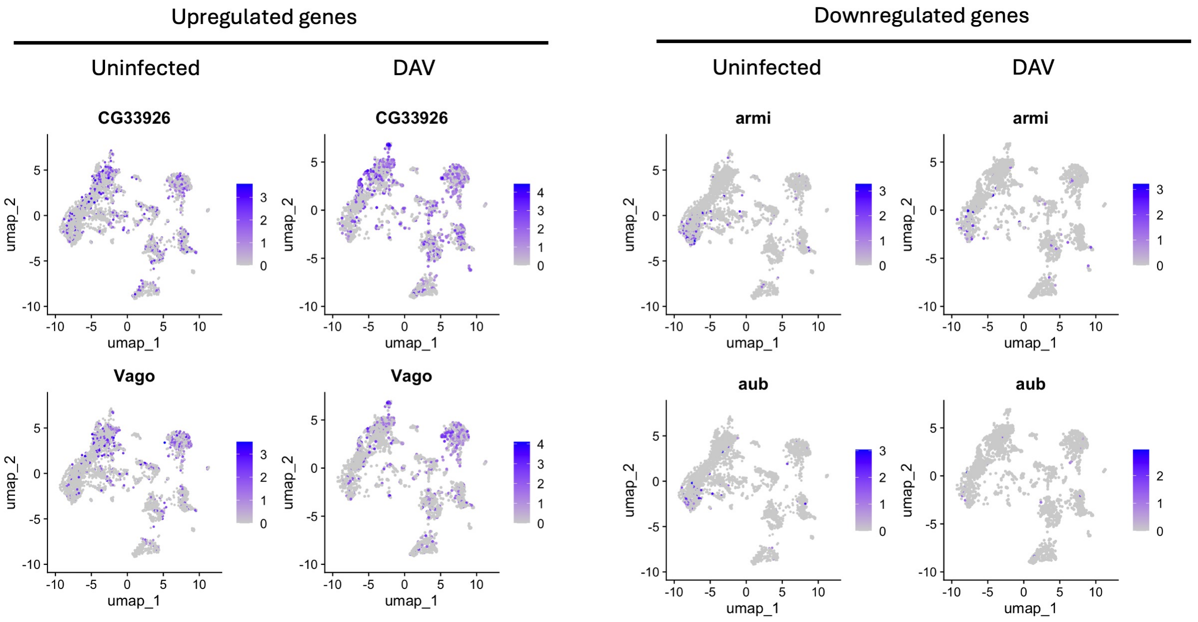


Figure S17: **UMAP-based comparison of differentially expressed gene (DEGs) expression between uninfected and DAV infected cells in mated flies.** Read counts are log normalized using Seurat LogNormalize method, where feature counts for each cell are divided by the total counts of that cell, multiplied by a scale factor of 10,000, and then taking natural log–transformed value using log1p. For CPM (counts per million) reference, log normalized counts can be converted to CPM using the formula $CPM = (e^{\text{LogNormalized count}} - 1) \times 100$. DAV= Drosophila A virus.

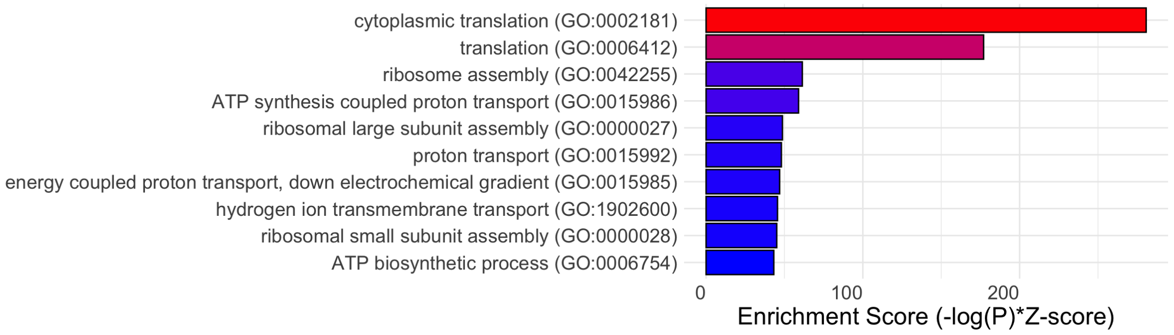


Figure S18: **Gene ontology (GO) enrichment analysis of differentially expressed genes in mated flies in Drosophila A virus infection.** Differentially expressed gene set was selected using a p-value threshold of $1 \times 10^{-5}$ for the GO analysis.

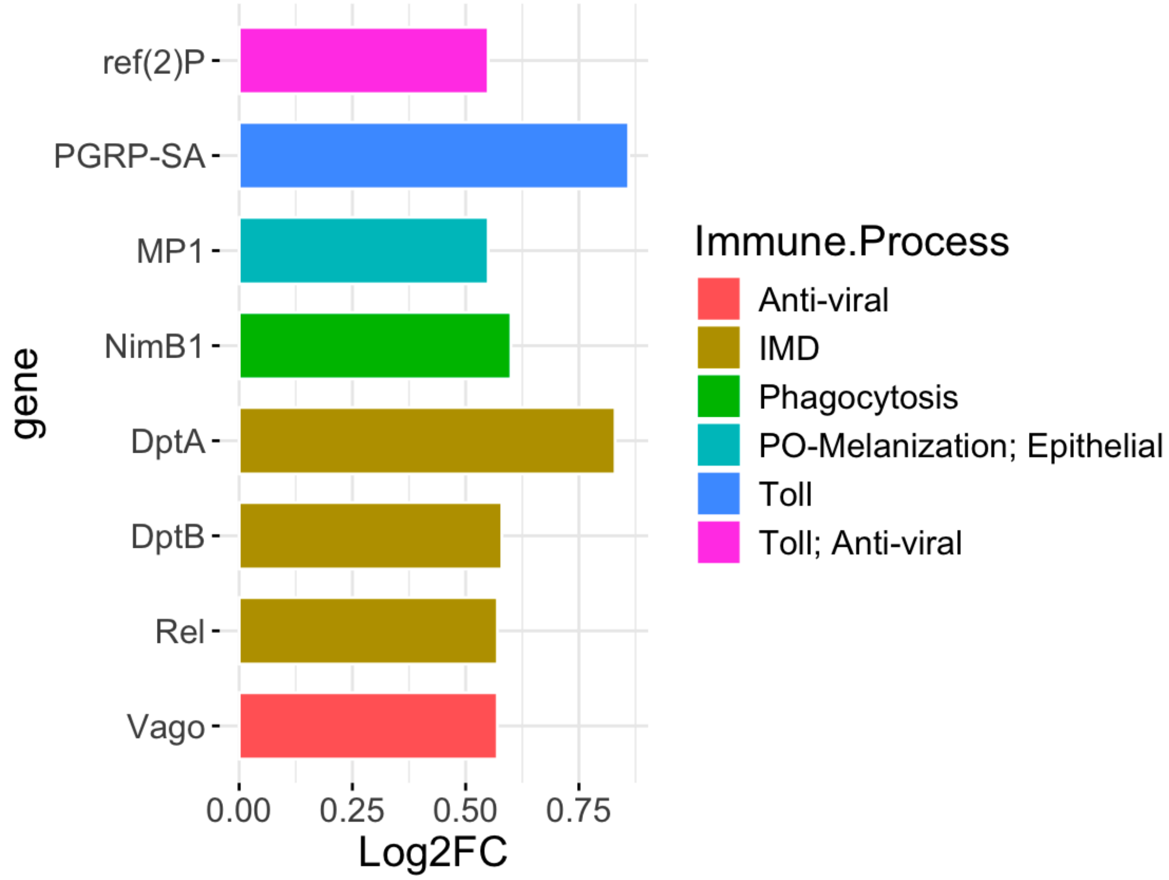


Figure S19: **Differentially expressed immune genes in mated flies during Drosophila A virus infection.** The gene set was selected using a p-adjusted (BH method) threshold of 0.05 and an absolute $\text{Log}_2$ fold change cutoff of 0.5. $\text{Log}_2\text{FC}$ = $\text{Log}_2$ fold change.

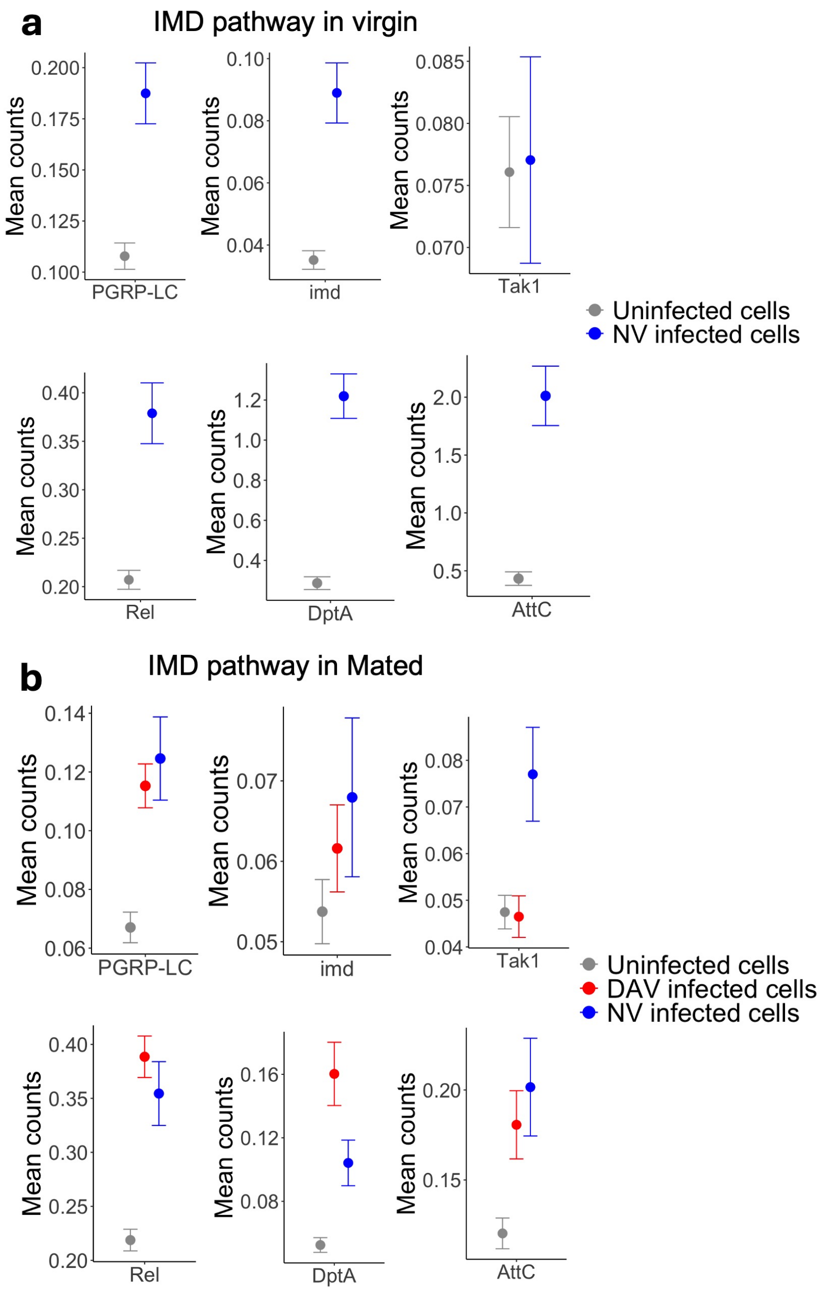


Figure S20: **Changes in IMD pathway genes during Nora and Drosophila A virus infection in virgin (a) and mated flies (b)**. IMD pathway is upregulated in association with viral infection. Read counts are log normalized using Seurat Log-Normalize method, where feature counts for each cell are divided by the total counts of that cell, multiplied by a scale factor of 10,000, and then taking natural log–transformed value using log1p. For CPM (counts per million) reference, log normalized counts can be converted to CPM using the formula $CPM = (e^{\text{LogNormalized count}} - 1) \times 100$. Uninfected cells are those with zero RNA reads for both Nora and Drosophila A viruses. DAV infected cells contain Drosophila A virus RNA reads (>0) but no Nora virus reads (0). NV infected cells contain Nora virus RNA reads (>0) but no Drosophila A virus reads (0). NV = Nora virus, DAV= Drosophila A virus.

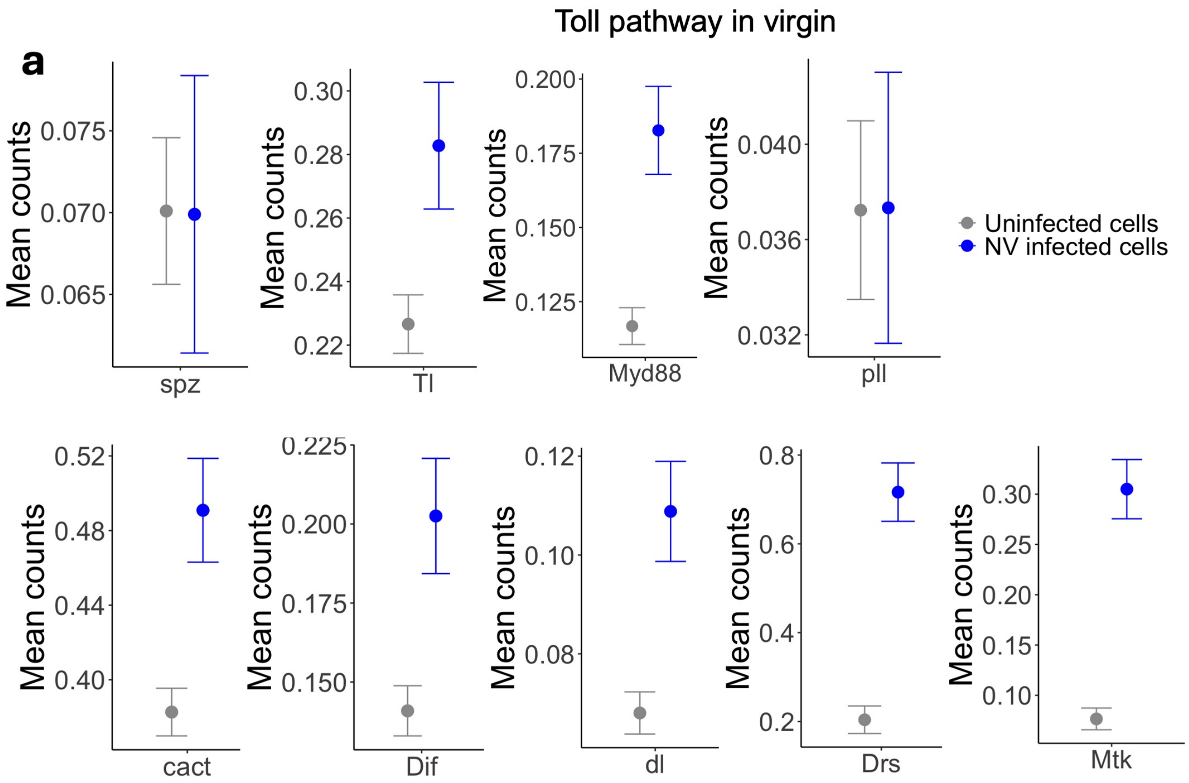


Figure S21: **Changes in Toll pathway genes during Nora virus infection in virgin files.** Toll pathway is upregulated in association with Nora virus infection. Read counts are log normalized using Seurat LogNormalize method, where feature counts for each cell are divided by the total counts of that cell, multiplied by a scale factor of 10,000, and then taking natural log–transformed value using log1p. For CPM (counts per million) reference, log normalized counts can be converted to CPM using the formula $CPM = (e^{\text{LogNormalized count}} - 1) \times 100$. Uninfected cells are those with zero RNA reads for Nora virus. NV infected cells contain Nora virus RNA reads (viral RNA >0). NV = Nora virus.

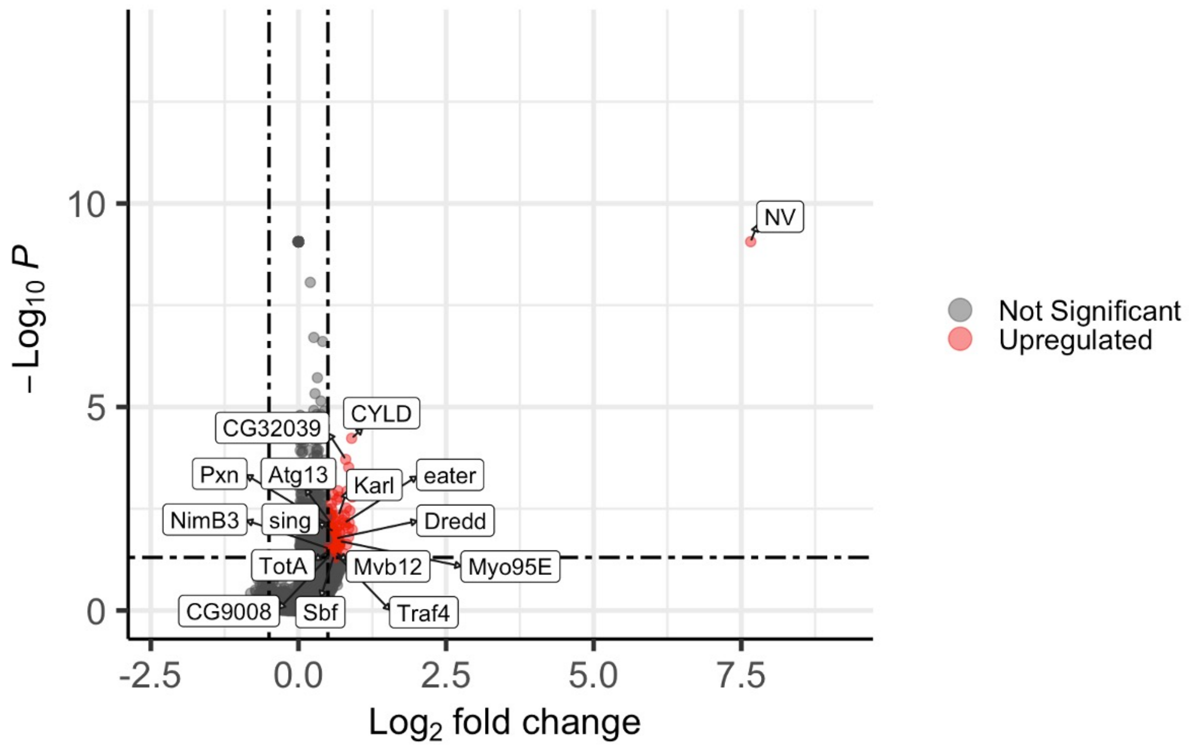


Figure S22: **Volcano plot of differentially expressed genes during Nora virus infection in mated flies.** In the mated condition, we concatenated replicates 1 and 2 (as in mated condition there were coinfection of Nora and Drosophila A virus). After concatenating, we compared Nora virus uninfected cells with Nora virus infected cells while controlling for cell types, Drosophila A virus infection, and replicates. (0.05/total number of genes) was used as *pvalue* threshold. Absolute value of 0.5 was used as threshold for $Log_2$ fold change ($Log_2$ FC). NV=Nora virus.

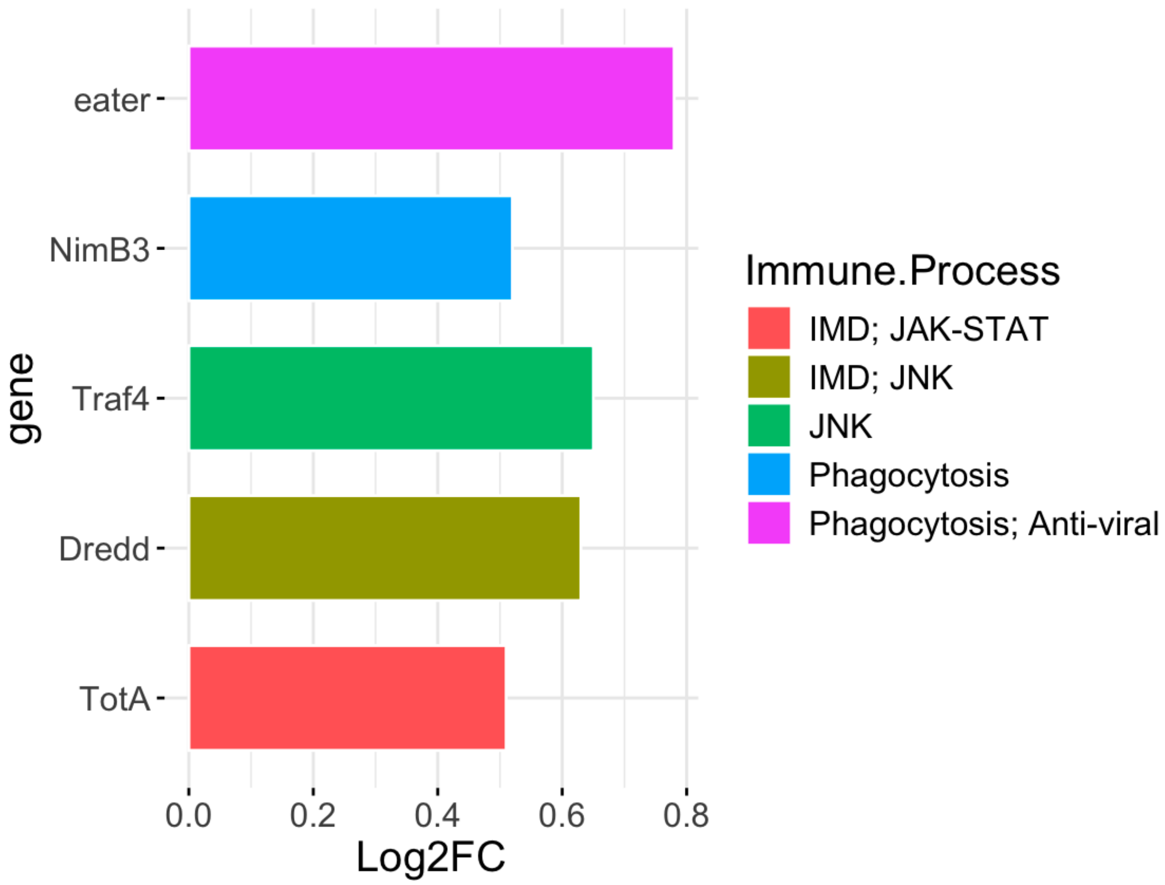


Figure S23: **Differentially expressed immune genes in mated flies in Nora virus infection.** The majority of the immune genes are upregulated in association with Nora virus infection. The gene set was selected using a p-adjusted (BH method) threshold of 0.05 and an absolute $\mathrm{Log_2}$ fold change cutoff of 0.5. $\mathrm{Log_2FC} = \mathrm{Log_2}$ fold change.

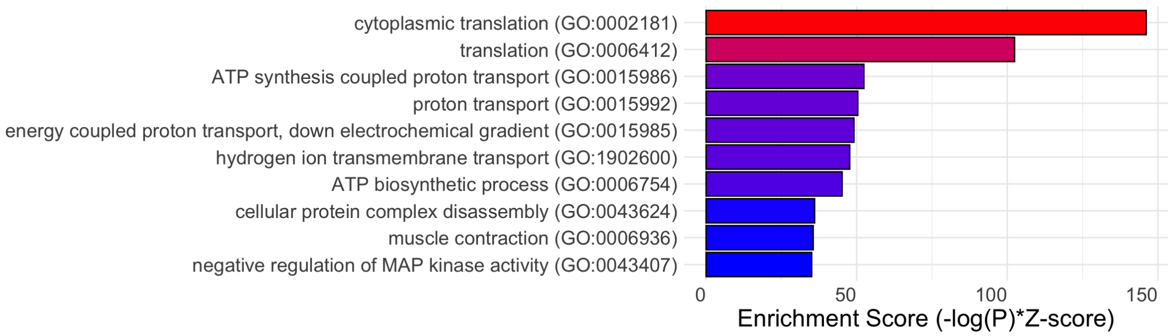


Figure S24: **Gene ontology (GO) enrichment analysis of differentially expressed genes in mated flies in Nora virus infection.** Differentially expressed gene set was selected using a p-value threshold of $1 \times 10^{-5}$ for the GO analysis.

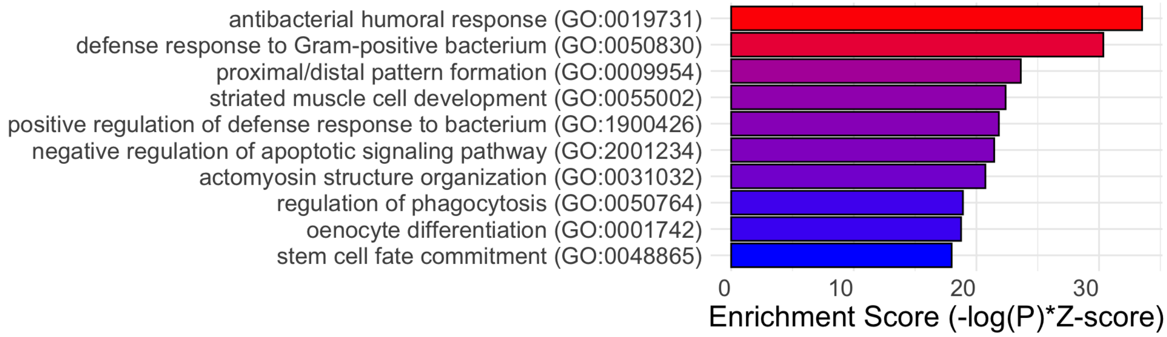


Figure S25: **Gene ontology (GO) enrichment analysis of interaction effect genes in virgin flies in Nora virus infection.** Here, the interaction effect means genes that change expression in viral infection based on cell types. The gene set was selected using a p-value threshold of $1 \times 10^{-5}$ for the GO analysis.

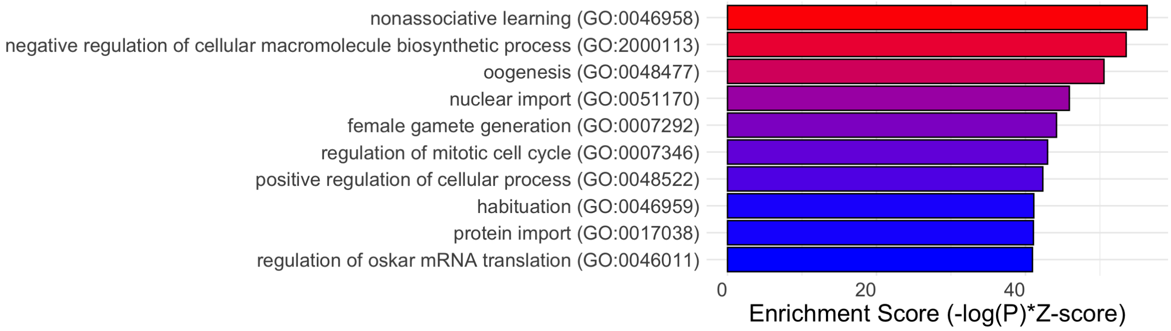


Figure S26: **Gene ontology (GO) enrichment analysis of interaction effect genes in mated flies in Drosophila A virus infection.** Here, the interaction effect means genes that change expression in viral infection based on cell types. The gene set was selected using a p-value threshold of $1 \times 10^{-5}$ for the GO analysis.

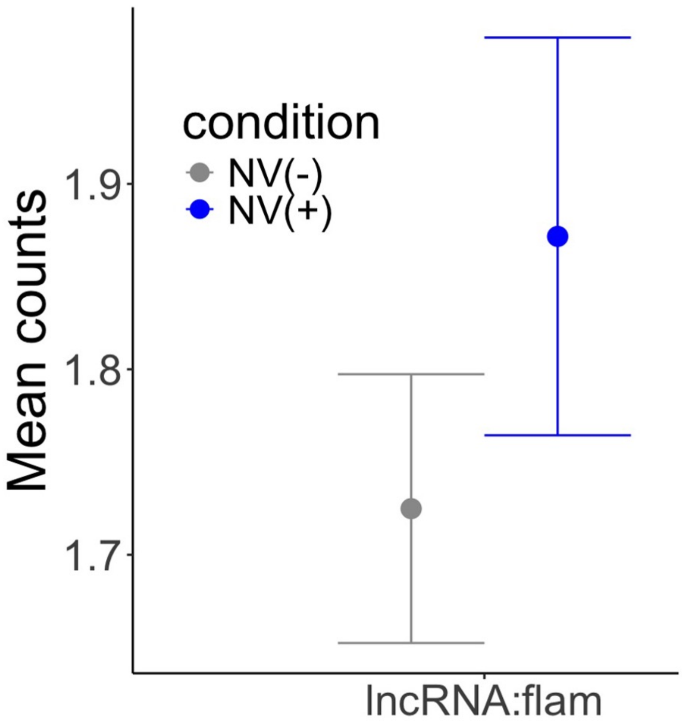


Figure S27: **Mean counts of flamenco (long noncoding RNA) gene in Nora virus infection in virgin flies**. Read counts are log normalized using Seurat LogNormalize method, where feature counts for each cell are divided by the total counts of that cell, multiplied by a scale factor of 10,000, and then taking natural log–transformed value using log1p. For CPM (counts per million) reference, log normalized counts can be converted to CPM using the formula $CPM = (e^{\text{LogNormalized count}} - 1) \times 100$. Uninfected cells are those with zero RNA reads for Nora virus. NV infected cells contain Nora virus RNA reads (viral RNA >0). NV = Nora virus.

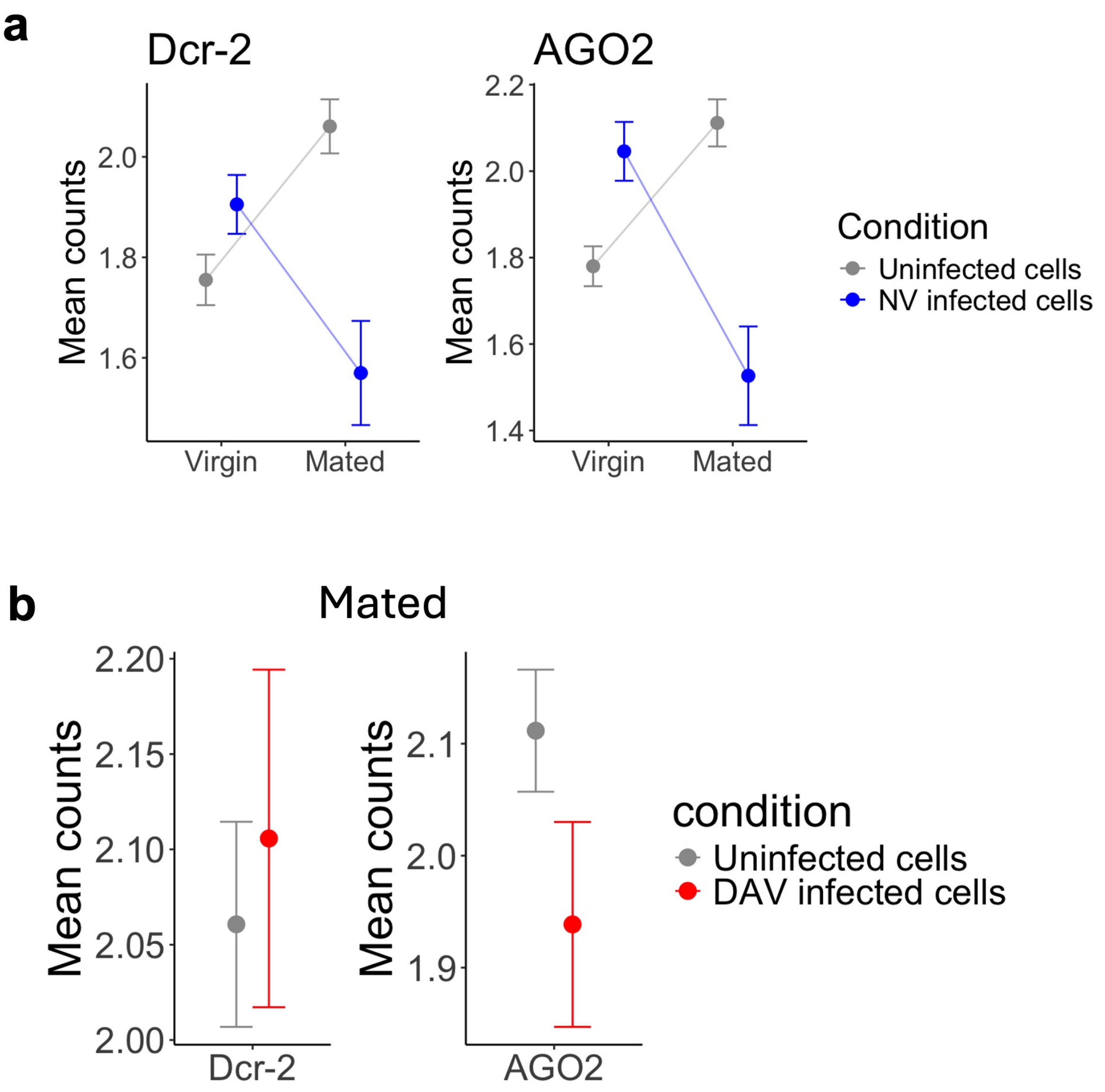


Figure S28: **Mean counts of RNAi pathway genes in Nora (a) and Drosophila A virus (b) infection.** Read counts are log normalized using Seurat LogNormalize method, where feature counts for each cell are divided by the total counts of that cell, multiplied by a scale factor of 10,000, and then taking natural log–transformed value using log1p. For CPM (counts per million) reference, log normalized counts can be converted to CPM using the formula $CPM = (e^{\text{LogNormalized count}} - 1) \times 100$. Uninfected cells are those with zero RNA reads for both Nora and Drosophila A viruses. DAV infected cells contain Drosophila A virus RNA reads (>0) but no Nora virus reads (0). NV infected cells contain Nora virus RNA reads (>0) but no Drosophila A virus reads (0). NV = Nora virus, DAV= Drosophila A virus.

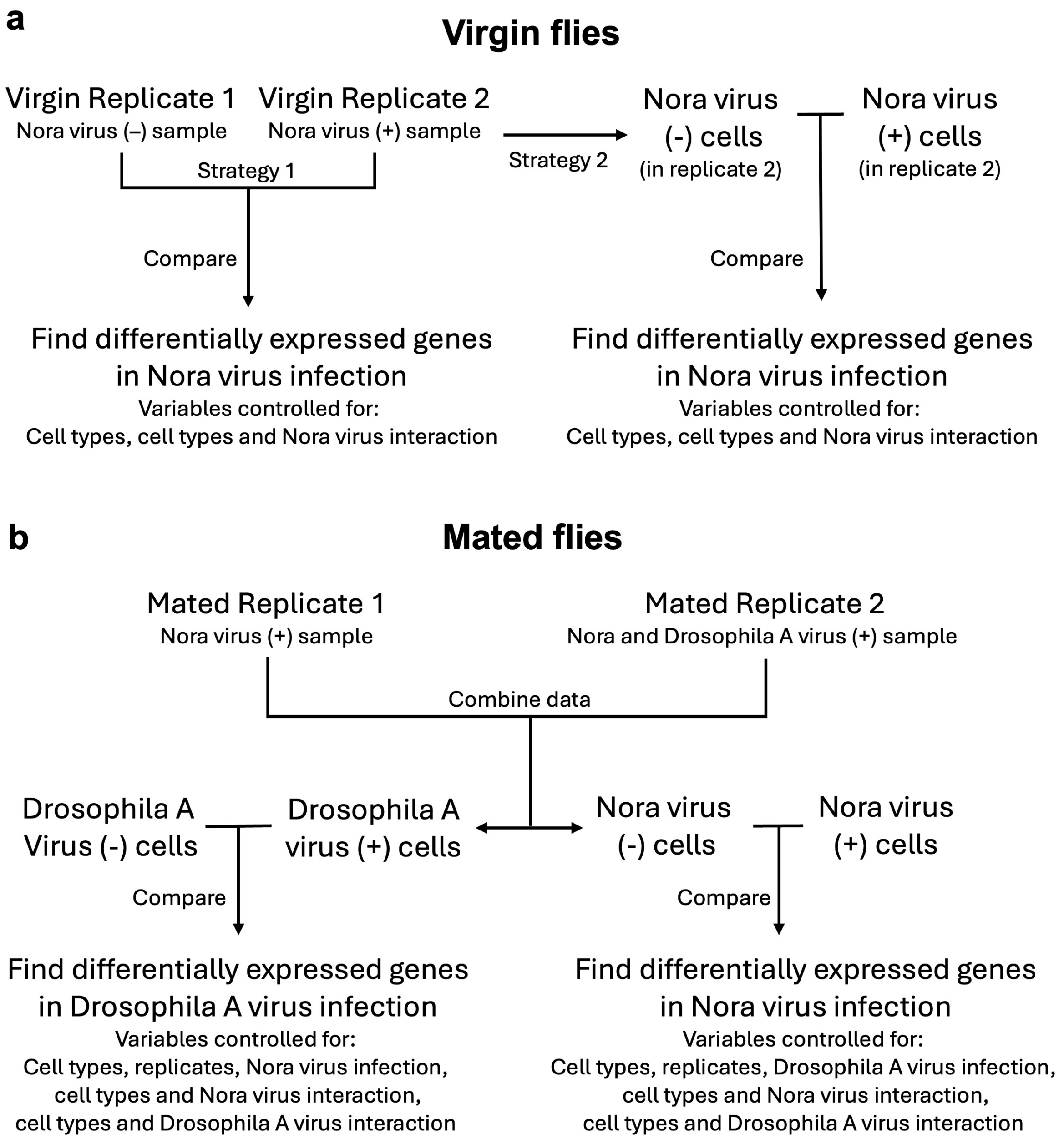


Figure S29: **The analysis methods to find differentially expressed genes in Nora (a) and Drosophila A virus (b) infection in virgin and mated flies.** This is a simplified schematic diagram. Refer to the method section to see the models used.
